# Three-color single-molecule imaging reveals conformational dynamics of dynein undergoing motility

**DOI:** 10.1101/2020.12.20.423706

**Authors:** Stefan Niekamp, Nico Stuurman, Nan Zhang, Ronald D. Vale

## Abstract

The motor protein dynein undergoes coordinated conformational changes of its domains during motility along microtubules. Previous single-molecule studies analyzed the motion of the AAA rings of the dynein homodimer, but not the distal microtubule binding domains (MTBD) that step along the track. Here, we simultaneously tracked two MTBDs and one AAA ring of a single dynein, as it undergoes hundreds of steps with nanometer precision using three-color imaging. We show that the AAA ring and the MTBDs do not always step simultaneously and can take different sized steps. This variability in the movement between AAA ring and MTBD results in an unexpectedly large number of conformational states of dynein during motility. Extracting data on conformational transition biases, we could accurately model dynein stepping in silico. Our results reveal that the flexibility between major dynein domains is critical for dynein motility.

## Introduction

The microtubule-based motor protein dynein belongs to the AAA+ (<u>A</u>TPases <u>A</u>ssociated with diverse cellular <u>A</u>ctivities) family of motors and is responsible for the majority of minus-end-directed motility along microtubules^1, 2^. Dyneins play key roles in many cellular processes and maintaining cellular architecture, including cargo transport, cilia motility, and the construction of the mitotic spindle^3–6^. Mutations or defects in cytoplasmic dynein are linked to several pathologies including cancers and neurological diseases^7, 8^.

Compared to kinesin^9,10^ and myosin^11,12^, cytoskeletal motors that have compact, globular motor domains, dynein is much larger and more complex with a size of ~1.4 MDa. Dynein is composed of two heavy chains and several associated polypeptide chains. The associated chains primarily bind to the N-terminal tail region to dimerize the heavy chains, regulate dynein’s function, and attach the motor to cargo^4,13,14^. The remaining two-thirds of the heavy chain constitute the motor domain, which is the driver of dynein motility^15^. The motor domain itself is divided into several domains - linker, AAA ring, stalk, and microtubule-binding domain (MTBD). The AAA ring consists of six different AAA domains that are linked together as an asymmetric hexameric ring (AAA1–AAA6) of which only AAA1-4 can bind ATP^16–19^. On top of the AAA ring lies the N-terminal linker that serves as a mechanical element and connects the motor domain to the N-terminal tail. The large catalytic AAA ring of dynein is separated from the small MTBD by a ~15 nm long, coiled-coil extending from AAA4 called the stalk^20–22^.

Upon ATP binding to AAA1, the motor domain releases from the microtubule and the linker undergoes the priming stroke (bending of the linker). During the priming stroke, the AAA ring has been observed to rotate relative to the linker and therewith bias the rebinding of the MTBD towards the microtubule minus end^1,23^. After ATP hydrolysis, the free MTBD rebinds to the microtubule while the linker undergoes the force-generating power stroke by straightening back to its initial conformation pulling the cargo with it^1,24–27^. Finally, with ADP release, the mechanochemical cycle can restart.

Initial dynein stepping experiments with a single fluorescent probe revealed that dynein, unlike kinesin, takes side- and backward steps^15^. In addition, dynein was shown to take variable step sizes, compared to kinesin, which only takes 8 nm steps^15,28^. Two-color single-molecule experiments revealed that the two AAA rings of dynein move in an uncoordinated manner, allowing one AAA ring to sometimes take multiple steps without any step of the other AAA ring^29,30^. In addition to the well-known hand-over-hand stepping of kinesin and myosin^28,31^, dynein can also move in an inch-worm fashion in which the leading AAA ring can step forward without movement of the trailing AAA ring, or in which the trailing AAA ring can step forward without passing the leading AAA ring^29,30^. Moreover, one active motor domain and an additional microtubule anchor are sufficient to achieve processive and directed motility^32^.

Prior single-molecule experiments followed the AAA ring, but not the MTBD that is actually stepping along the microtubule track. Since the AAA ring and the MTBD are separated by the ~15 nm stalk, which can adopt different angles with respect to the MTBD^2,33,34^, the position and stepping of the AAA ring may not reflect that of the MTBD. Thus, to have an accurate understanding of dynein stepping, it is important to directly measure the position of the MTBDs relative to the microtubule track and to measure the position of the MTBDs relative to the AAA rings.

Here, we developed a three-color single-molecule microscopy assay that allows simultaneous tracking of the movement of one AAA ring and two MTBDs for the first time. In addition to extending existing nanometer accuracy distance measurement and image registration, we also utilized ~6 nm small fluorescent probes (DNA FluoroCubes) that are ~50-fold more photostable than organic dyes^35^, allowing many dynein steps to be measured without photobleaching. Using these technical advances, we show that the AAA ring and MTBD sometimes step at different times and take differently sized steps, which gives rise to a large variety of conformations that dynein can adopt as it walks along the track. The transition probabilities between conformations derived from our data are sufficient to recapitulate directed dynein motility using Monte Carlo simulations. Taken together, we conclude that dynein can adopt many conformations, including some previously undescribed ones, and that the AAA ring and MTBD exhibit different stepping behaviors.

## Results

### Development of a three-color dynein imaging system

To determine how the AAA ring and MTBD move relative to each other while dynein is stepping along microtubules, we tracked the stepping of a three-color labeled dynein in which one AAA ring and two MTBDs were labeled with three differently colored fluorophores (**Fig. 1 a**). To allow accurate tracking of all three colors with respect to one another with 1 nm resolution, we extended our previously developed two-color image registration routine^36^ to three colors (**Supplementary Fig. 1 a-d**). To validate this approach, we imaged three differently colored dyes placed at well-defined distances on a DNA-origami nanoruler^37,38^ and found that the expected distances among the three dyes were recovered with 1 nm accuracy (**Supplementary Fig. 1 e-g**).

**Figure 1.**
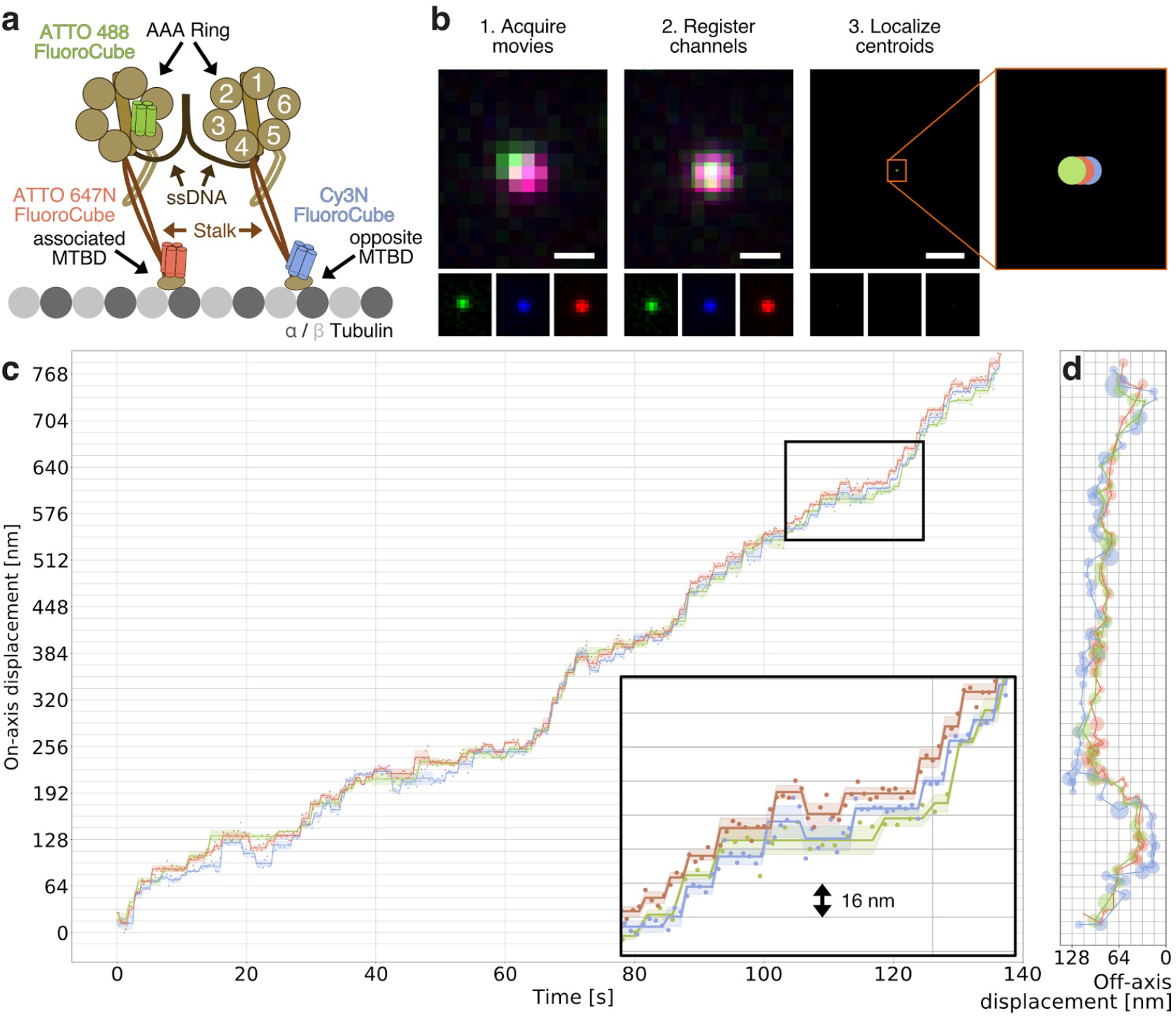
Three-color stepping trace of dynein. (**a**) Schematic of design of three-color dynein. Each of the two motor domains of dynein is labeled individually and dimerized using reverse-complementary single-stranded DNA (black, attachment via SNAP-tag^39,41^). The MTBD of each motor domain and one of the two AAA rings are labeled with FluoroCubes^35^. For one motor domain, the AAA ring is labeled with a six dye ATTO 488 FluoroCube (green, attachment via HALO-tag^40^) and the MTBD (termed associated MTBD) is labeled with a six dye ATTO 647N FluoroCube (red, attachment via YBBR-tag^41^). For the other motor domain only the MTBD is labeled (termed opposite MTBD) with a six dye Cy3N FluoroCube (blue, attachment via YBBR-tag^41^). More details about construct design and labeling can be found in **Materials and Methods**. (**b**) Workflow to collect three-color dynein stepping data. The top micrographs show a merge of all three colors while the bottom row shows each color separately. Scale bar is 500 nm. More details on data collection can be found in **Materials and Methods**. (**c**) Raw stepping data with position along the on-axis versus time of a three-color dynein heterodimer (colored dots) with detected steps (colored lines). The opaque lines show the standard deviation along the on-axis for each step. Insert is a magnified view of the area in the black box on the bottom left. (**d**) The same trace as in c but in xy space. The colored circles show the fitted position, with the radius corresponding to the standard error of the mean of the combined on- and off-axis.

To create a three-color labeled dynein dimer in which one AAA ring and two MTBDs are fluorescently labeled, we used the well-studied, truncated yeast cytoplasmic dynein^15^ and added a N-terminal SNAP-tag^39^, a C-terminal HALO-tag^40^, and an internal YBBR-tag^41^. In this design, the HALO-tag is positioned on top of the AAA ring and the YBBR-tag is placed in a flexible loop of the MTBD, enabling us to label both the AAA ring and the MTBD on the same motor domain. For one monomeric motor domain, we labeled the HALO-tag with a six-dye ATTO 488 FluoroCube^35^ and the YBBR-tagged MTBD with a six-dye ATTO 674N FluoroCube. For another monomeric motor domain, we only labeled the YBBR-tagged MTBD with a six dye Cy3N FluoroCube (**Fig. 1 a**). To join the two-labeled monomers together into a dimer, we separately labeled the N-terminal SNAP-tag on the monomeric motor domains with reverse-complementary single-stranded DNAs. When combined together, the hybridization of the single-stranded DNAs created a dimeric motor as previously described^29^. This three-color FluoroCube labeled dynein had a similar velocity and processivity as a GFP-tagged wild-type dynein (**Supplementary Fig. 2 a-c, Supplementary Movie 1**), suggesting that FluoroCube labeling did not perturb dynein function.

When we compared a three-color dynein labeled with conventional organic dyes to a dynein labeled with FluoroCubes, we found that 4% of the conventional dye-labeled dynein had a signal in all three channels after 50 frames while 75% of FluoroCube-labeled dyneins emitted signals in all three channels (**Supplementary Fig. 2 f, g**). Moreover, the FluoroCube-labeled dynein yielded more precise localizations compared to conventional dyes for the same exposure time of 110 ms (2.4 nm for a Cy3N FluoroCube compared to 7.2 nm for a single conventional dye of the same color; **Supplementary Fig. 2 d, e**). In summary, without using FluoroCubes, the tracking of all three domains simultaneously with high resolution would not have been feasible.

### Step size analysis of a three-color dynein

We tracked the stepping of three-color-labeled dyneins at low ATP concentration (3 μM) along microtubules to resolve individual steps of all three domains at high spatiotemporal resolution (**Fig. 1 b-d, Supplementary Movie 2**). In order to enable a fast acquisition of 330 ms with minimal dead time, we optimized the acquisition sequence (see Materials and Methods, **Supplementary Fig. 3**) and resolved 4,500 steps from >54 dynein molecules moving along microtubules (**Fig. 1 c, d**). Using this dataset, we found a similar average step size and percentage of forward steps for the AAA ring when compared to previous studies^29,30^ (**Fig. 2, Supplementary Fig. 4**). In addition, we were able to analyze the stepping behaviour of the MTBDs for the first time (**Fig. 2**). We observed that the MTBDs tend to not pass each other, resulting in inch-worm stepping behaviour (**Supplementary Fig. 4**), as previously described for the two-color-labeled AAA rings^29,30^. Also in agreement with previous studies of AAA rings^29,30^, the trailing MTBD was more likely to take the next step compared to the leading MTBD. However, we found significant quantitative differences between the stepping of MTBDs and AAA rings. For example, the AAA ring, on average, took slightly larger forward steps (22.2 nm) compared to the MTBD (18.8 nm for both MTBDs combined). Moreover, the AAA ring took fewer backward steps (14%) compared to the MTBD (21% and 19% for the blue and red labeled MTBD, respectively). However, as expected, the step size distributions of both MTBDs were identical (**Fig. 2 c, h**). The off-axis step sizes of the AAA ring and MTBD were also very similar (**Fig. 2 d-h, Supplementary Fig. 4 a**). Moreover, we obtained similar results for three-color dynein moving along axonemes compared to microtubules (**Supplementary Fig. 3**). Thus, these results show that the two MTBDs exhibit identical movements, while the AAA ring and the MTBD do not, indicating more complicated movements rather than a simple rigid body translation of the entire motor domain.

**Figure 2.**
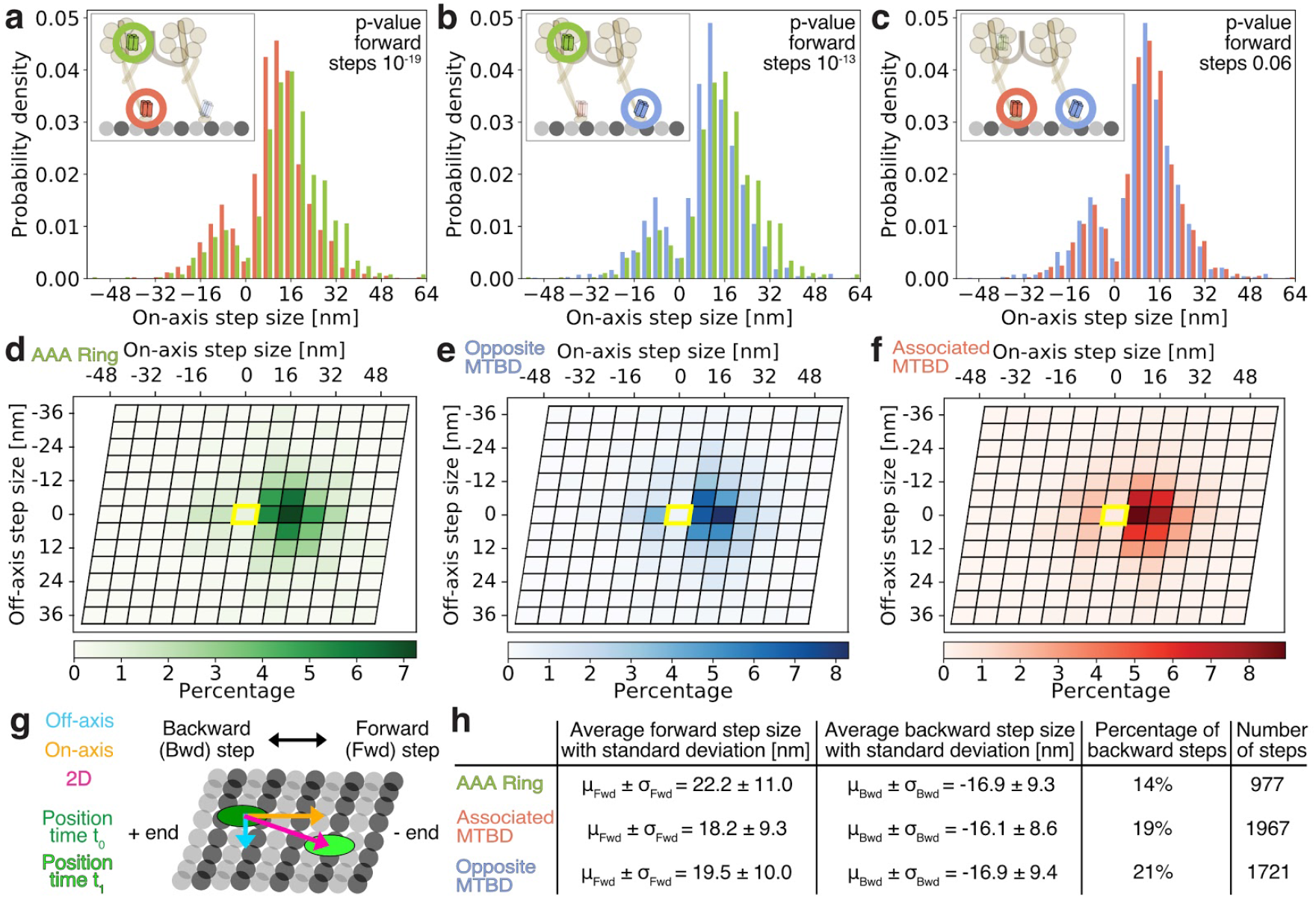
Two-dimensional analysis of AAA ring and MTBD stepping. (**a**) Histogram of on-axis step sizes of dynein’s AAA ring (green) and the MTBD (red) of the same motor domain (associated MTBD). (**b**) Histogram of on-axis step sizes of dynein’s AAA ring (green) and the MTBD (blue) on the opposite motor domain. (**c**) Histogram of on-axis step sizes of dynein’s MTBDs (blue and red). (**a-c**) Box in the top left shows a schematic which domains of dynein were analyzed. The p-values were calculated with a two-tailed t-Test. (**d-f**) Same data as in **a-c** shown as a heatmap of the on-and off-axis step sizes of (**d**) dynein’s AAA ring (green), (**e**) the opposite MTBD (blue), and (**f**) the associated MTBD (red) mapped on a microtubule lattice. The microtubule lattice is based on a 13 protofilament microtubule. Here, each parallelogram represents a tubulin dimer consisting of one copy of and tubulin. The yellow parallelogram represents the tubulin dimer at which the domain was located prior to the step. (**g**) Microtubule lattice (grey circles) with plus and minus ends and the definition of forward and backward as well as 2D, on- and off-axis steps. (**h**) Table summarizing properties of step size distributions shown above for dynein’s AAA ring (green), the associated MTBD (red), and the opposite MTBD (blue).

### Independent stepping of the MTBD and the AAA ring

Given the difference in on-axis step sizes and percentages of forward and backward steps of the AAA ring and MTBD, we next examined the timing of the steps of these domains (**Fig. 3 a**). When the MTBD takes a short step (4-12 nm; centered around the dimension of a tubulin subunit (8 nm)), the AAA ring on the same motor domain displays an evident simultaneous step only ~60% of the time. Thus, not every step of a MTBD results in the relocation of the AAA ring on the same motor domain. However, when the MTBD stepped by distances of >20 nm (corresponding to distances of approximately three or more tubulin subunits), the probability of simultaneous stepping of the AAA ring increased to >90% (**Fig. 3 b**). Collectively, these results might be explained by flexibility between these domains; when the MTBD takes a short step, the stalk can adjust its angle relative to the AAA ring, resulting in little axial displacement of the AAA ring. However, longer MTBD steps can only be accommodated by a displacement of the AAA ring along the microtubule axis.

**Figure 3.**
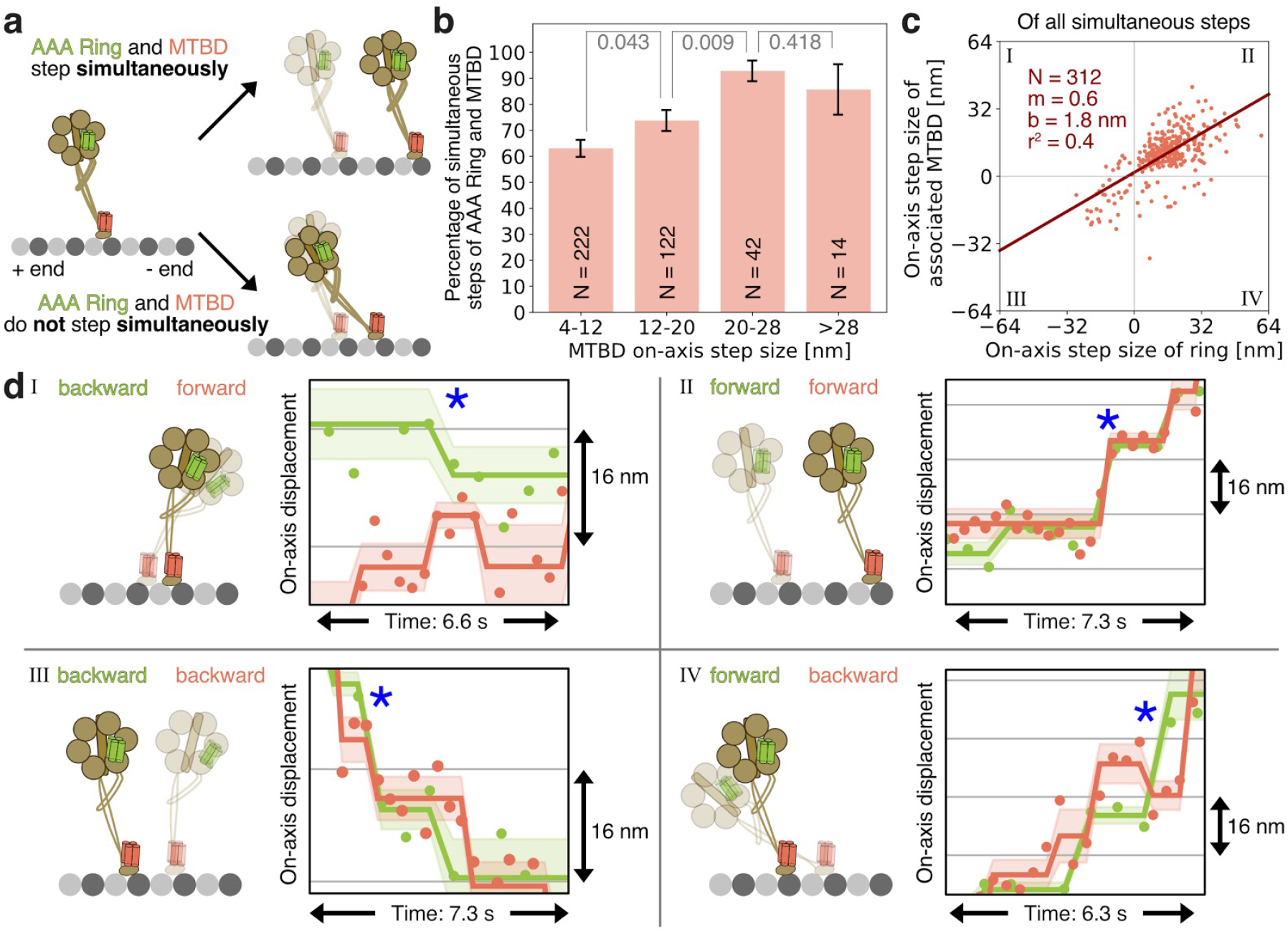
Independent stepping of AAA ring and MTBD on the same motor domain. (**a**) The AAA ring and MTBD on the same motor domain can either step simultaneously (top: both domains move along the on-axis) or not step simultaneously (bottom: only MTBD moves while the AAA ring remains at the same on-axis position). (**b**) Histogram showing how often the AAA ring steps at the same time as the associated MTBD (red) as a function of the MTBD on-axis step size. N refers to the total number of steps for each condition. The error bars show the bootstrapped standard error of the mean. The p-values (grey) were calculated with a two-tailed z-Test. (**c**) Correlation of on-axis step sizes of the AAA ring and the associated MTBD (red) when they step at the same time. Each dot represents a single step. Red line shows linear fit. N is the sample size. m is the slope. b is the y-intercept. r^2^ is r squared value. (**d**) Example on-axis traces for each of the four quadrants (I, II, III, IV) defined in **c**, accompanied by a diagrammatic representation where the initial position is shown with decreased opacity. The blue stars indicate the time at which the AAA ring and MTBD moved simultaneously. All traces are raw stepping data with position along the on-axis versus time of a three-color dynein heterodimer (colored dots) with detected steps (colored lines). The opaque lines show the standard deviation along the on-axis for each step. Note, the blue channel was removed for clarity.

We also examined how often the MTBD on one motor domain steps at the same time as the AAA ring on the other motor domain and found a lower probability for them to step simultaneously compared to MTBD and AAA ring of the same motor domain (**Supplementary Fig. 5 a-c**). Thus, a step of a MTBD of one motor domain also does not necessarily result in the axial displacement of the AAA ring on the other motor domain.

Next, we focused our analysis on the direction and dimensions of the simultaneous steps of AAA ring and MTBD of the same motor domain (**Fig. 3 b**). We found that the relative step sizes and directions were not always the same (**Fig. 3 c, d**). Although in most cases both domains stepped forward (**Fig. 3 c, d** quadrants II), we observed cases in which the domains took steps in opposite directions: e.g. the MTBD takes a short backward step while the AAA ring moves slightly forward (**Fig. 3 c, d** quadrant IV) and vice versa (**Fig. 3 c, d** quadrant I). The least likely event is for the MTBD to step forward and the AAA ring to move backwards ((**Fig. 3 c, d** quadrant I). In conclusion, AAA ring and MTBD do not necessarily advance along the track at the same time and over the same distance, further supporting the idea that they are not rigidly connected bodies.

### Relative positions of the AAA ring and the MTBD

To investigate how the AAA ring and the MTBD move relative to each other, we examined the relative positions between green-labeled AAA ring and red-labeled MTBD of the same motor domain (**Fig. 4 a**). On average, the MTBD was positioned in front of the AAA ring (closer to the microtubule minus end) (**Fig. 4 b**). This finding is consistent with static electron microscopy data of dynein bound to microtubules^2,34^. However, there was no preferential position of the two MTBDs or between the MTBD and AAA ring along the off-axis of the microtubule (**Supplementary Fig. 5 d-i**).

**Figure 4.**
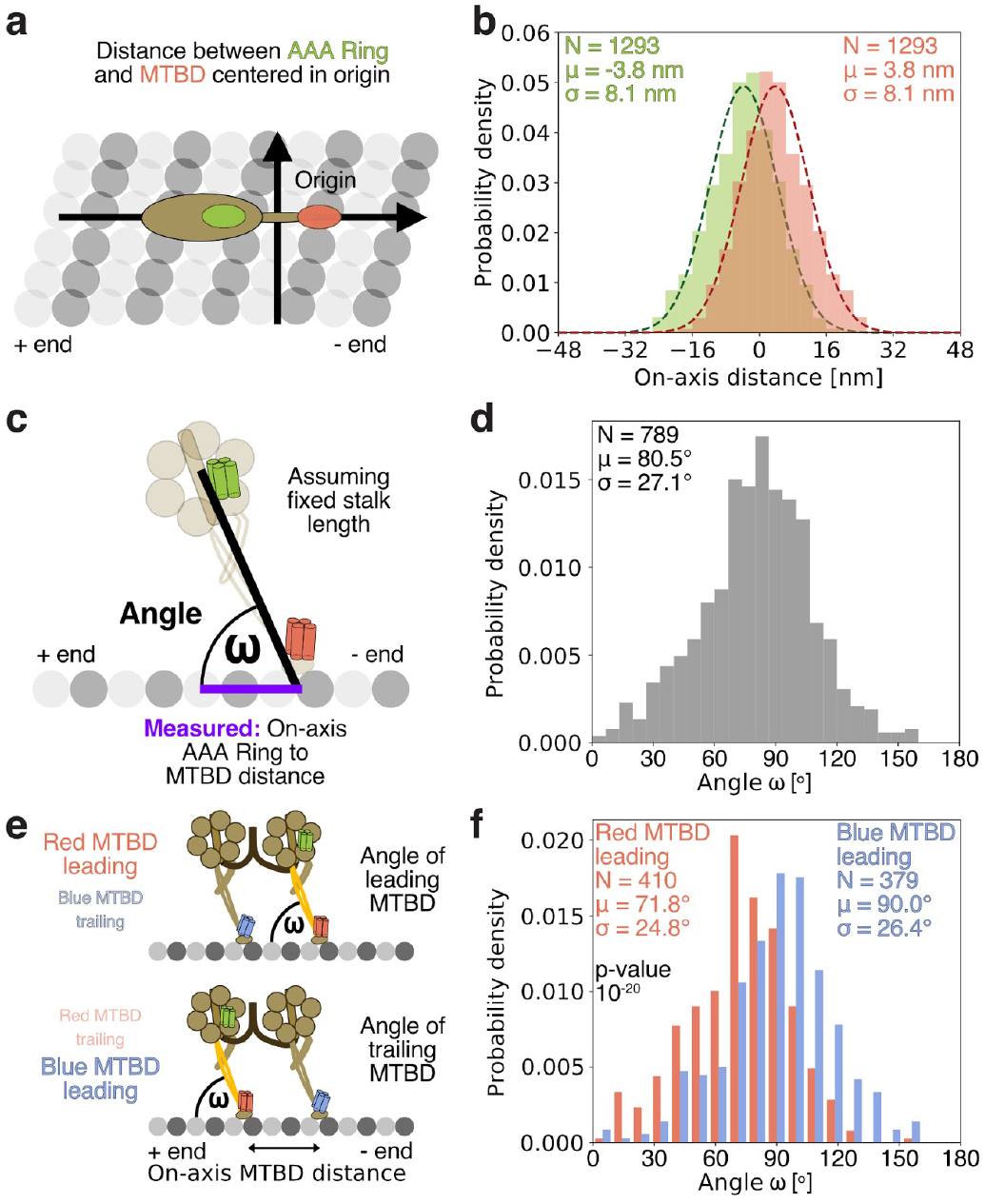
Relative movement of AAA ring and MTBD on the same motor domain. (**a**) Distance between AAA ring and MTBD of same motor domain. Here, the centroid position of AAA ring and MTBD are fixed in the origin and the distance of both domains relative to the centroid is measured. (**b**) Histogram of on-axis distances between AAA ring (green) and the MTBD (red) of the same motor domain. Sample size (N), average distance (μ) and its standard deviation (σ) is given. (**c**) Schematic showing the definition of the angle ω between stalk and microtubule on-axis. Note, the angle is only calculated for the motor domain for which the AAA ring (green) and the MTBD (red) are both labeled. To calculate the angle ω we used the measured on-axis distance between the AAA ring and MTBD (purple line) and the fixed known distance from the MTBD to the center of the AAA ring (black line). (**d**) Histogram of stalk-microtubule angles ω. (**e**) Schematic showing definition of leading and trailing MTBD. Top: Angle measurement for leading MTBD (red MTBD leading). Bottom: Angle measurement for trailing MTBD (blue MTBD leading). (**f**) Histogram of stalk-microtubule angles ω if either the motor domain is leading (red MTBD leading) or trailing (blue MTBD leading). The p-value was calculated with a two-tailed t-Test. (**d, f**) Sample size (N), average distance (μ) and its standard deviation (σ) is given.

Since we observed a large spread of distances between AAA ring and MTBD on the same motor domain, we next asked how this influences the angle between stalk and microtubule. To calculate the stalk-microtubule angle ω, we used the on-axis distance between the AAA ring and MTBD of the same motor domain and assumed a fixed length of the dynein stalk (**Fig. 4 c**). We found a wide distribution of angles averaging at 80.5° (**Fig. 4 d**), further supporting the idea of a high flexibility within the dynein motor domain.

Next, we asked whether the stalk-microtubule angle ω is different for the leading and trailing motor domains (**Fig. 4 e**). Comparing the average angle for both cases, we found that the angle ω for the leading motor domain (red MTBD leading; 71.8°) was significantly smaller than for the trailing motor domain (red MTBD trailing; 90.0°) (**Fig. 4 f**). We also calculated the stalk-microtubule angle ω as a function of inter-MTBD on-axis distance and found that the angle of the trailing motor domain increases with increasing separation of the two MTBDs while the angle of the leading motor domain decreases (**Supplementary Fig. 5 j-l**). Taken together, these data reveal flexibility between the AAA ring and MTBD on the same motor domain and that the leading motor domain typically adopts a more acute stalk-microtubule angle compared with the trailing motor domain.

### Dynein adopts a large variety of conformations

Three-color imaging enabled us to determine the relative positions of the AAA ring and the two MTBDs along the on-axis of the microtubule. Permuting through all possible orders of three colors leads to a total of six different domain orderings, each of which can be associated with four potential conformational states of dynein (**Fig. 5 a, Supplementary Fig. 6 a, b**). To determine the relative frequency of domain orders, we quantified how often each of the six domain orders occurs during all stepping traces. We found that dynein can adopt all six domain orders to a varying degree (**Fig. 5 b**). The two most common domain orders were the ones in which both MTBDs are leading the green-labeled AAA ring (51% combined), followed by the two domain orders in which the AAA ring is positioned between both MTBDs (32% combined), and followed by the two domain orders in which the AAA ring is leading (17% combined). Interestingly, based on previous studies^1^ we would not have predicted the two domain orders in which the green-labeled AAA ring is leading. Together, the observation of a large variety of domain orders suggests that dynein can adopt a large variety of conformational states during motility (**Supplementary Fig. 6 a, b**).

**Figure 5.**
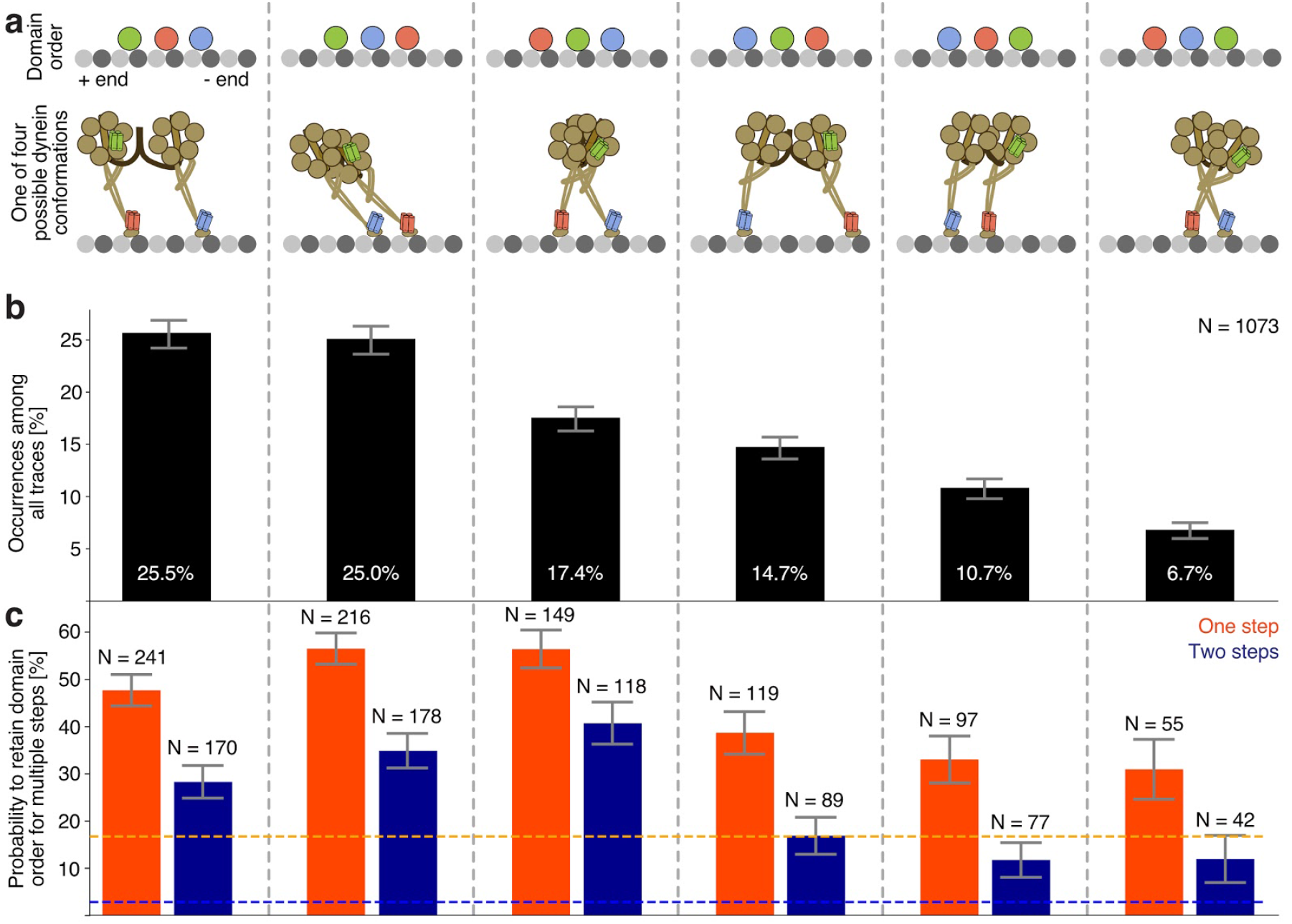
Frequency of domain orders of AAA Ring and both MTBDs. (**a**) Schematic of all six possible domain orders along the microtubule on-axis. Top: small, grey circles show tubulin while the larger green, blue, and red circles represent the AAA ring, the opposite MTBD, and the associated MTBD, respectively. For instance, to be classified as the very left domain order, the opposite MTBD (blue) has to be closest to the microtubule minus end, followed by the associated MTBD (red) and followed by the AAA ring (green). Note that the absolute distance between domains is irrelevant. Bottom: One of the four possible dynein conformations based on the three color domain orders. Other possible conformations are shown in **Supplementary Figure 6**. (**b**) Histogram of occurrence of each of the six possible domain orders with sample size N. The error bars show the bootstrapped standard error of the mean. (**c**) Probability to retain domain order after one (orange) or two (blue) steps. Here, a step refers to the movement of at least one of the three domains. The orange and blue dotted lines indicate the probability to retain the domain order after one (orange) or two (blue) steps if transitions were random. The sample size N refers to the total number of all domain order transitions that occurred after the motor took a step out of its current domain order. The error bars show the bootstrapped standard error of the mean. A more detailed analysis of transitions between domain orders is given in **Supplementary Figure 5**.

We next asked how frequently dynein transitions to a new domain order after a step occurred. If conformational states were random, we would have expected that 1/6 (~17%) will remain in the initial domain order after one step and only 1/36 (< 3%) will still have the same domain order after the first and second step. However, we found that dynein tends to remain in its same domain order after a step (**Fig. 5 c, Supplementary Fig. 5 m**), although some states are more persistent than others. For example, the two domain orders in which the green-labeled AAA ring is leading are the least stable and more likely to transition to other domain orders in which the MTBDs are leading. This observation of a persistence of domain orders agrees with previous observations in which the AAA rings of dynein were reported to infrequently pass each other^29^. We also measured how often a step of any of the labeled domains was followed by a step of the same or another domain. The most common outcome was that one MTBD moves after the other (**Supplementary Fig. 5 n**), which is again consistent with the observation that the two dynein AAA rings tend to step in an alternating fashion^29,30^. In summary, dynein’s AAA ring and the two MTBDs often move in an alternating fashion and are less likely to pass each other, resulting in a persistence of a given domain order over multiple steps.

### Simulation of dynein motility

The observation of six three-color domain orders shows that dynein can adopt a large variety of conformations when moving along microtubules. However, we lack information on the location of the second AAA ring. Thus, we turned to Monte Carlo simulations to obtain more insights into dynein conformations during motility. Using our experimental data as input, we simulated the stepping of both AAA rings and both MTBDs along microtubules (**Fig. 6 a, Supplementary Movie 3**) by assigning probabilities to step sizes, stepping directions, and likelihoods of what kind of step will follow after another, as described in Materials and Methods. We also applied a few rules that are based on our data of the three-color dynein (**Supplementary Movies 4-9**). Specifically, (1) an on-axis distance-dependent bias to take more forward than backward steps (**Fig. 2**), (2) a distance-dependent bias to close the gap between the motor domains along the on- and off-axis when taking a step (**Supplementary Fig. 4**), (3) a higher probability for the trailing domain instead of the leading domain to take the next step (**Supplementary Fig. 4**), (4) a bias towards alternating stepping behavior (**Supplementary Fig. 4**), and (5) the relative movement between AAA ring and MTBD (changes in angle ω) (**Fig. 4**). However, we did not enforce specific distances between the two motor domains by setting cutoffs for on- and off-axis distance; in other words, the motor domains were not constrained by a connecting tether and stepped independently (**Fig. 6 a**) according to the rules set above.

**Figure 6.**
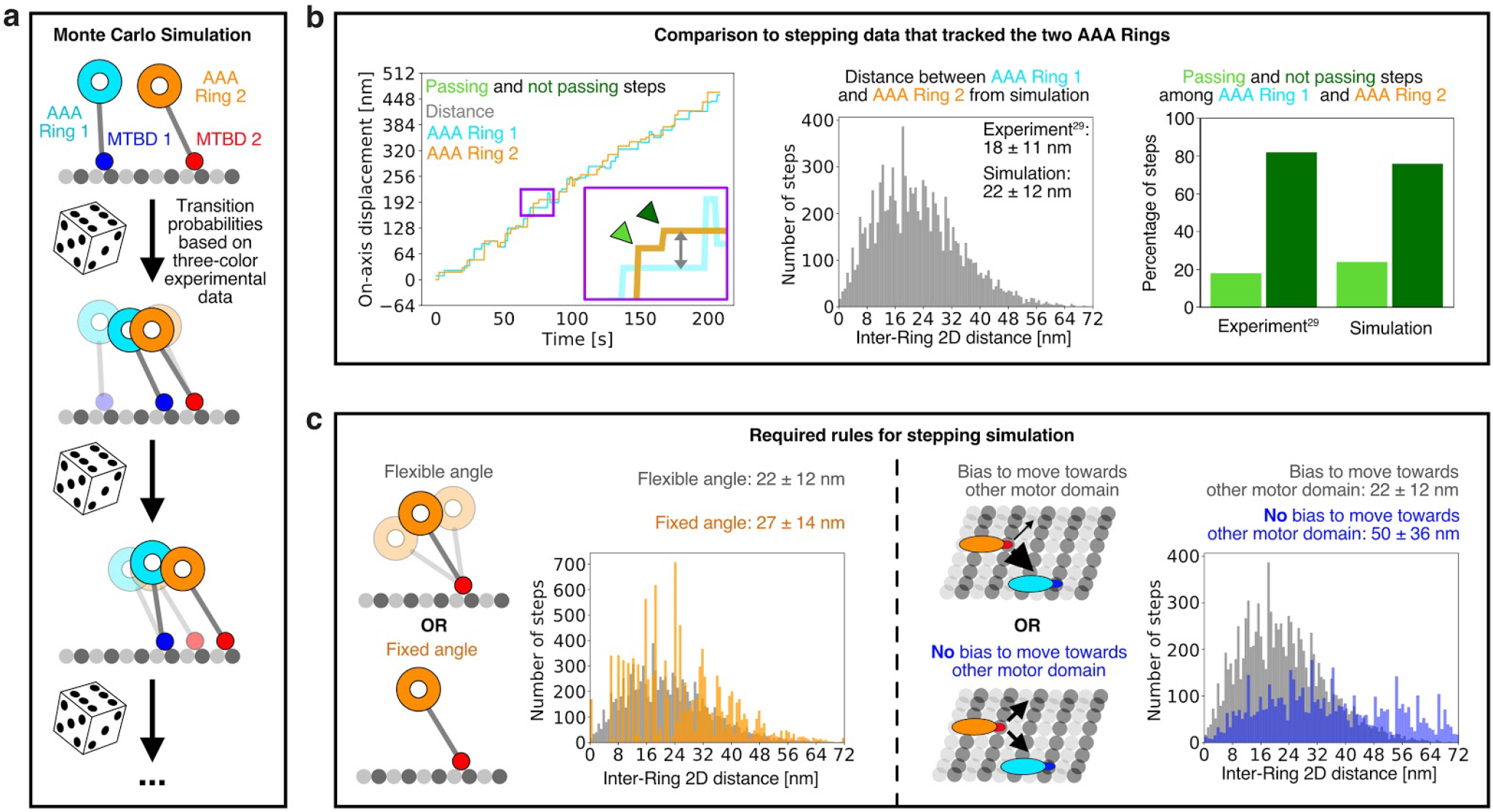
Monte Carlo simulation of dynein motility. (**a**) Using the three-color experimental data as input the positions of both MTBDs and both AAA rings can be simulated using Monte Carlo simulation. (**b**) Comparison of Monte Carlo simulated dynein motility to previously published experimental data for which both AAA Rings were tracked^29,30^. Left: Example stepping traces from Monte Carlo simulation of both AAA rings. Middle: Histogram of 2D distance between both AAA rings compared to previously measured inter-ring 2D distance^29,30^. Right: Passing and not passing steps among the two AAA rings based on simulations compared to previously measured data^29^. (**c**) By running Monte Carlo simulations the importance of different rules on dynein motility can be evaluated. Here, the influence of two rules on the 2D distance between both AAA rings was tested. (1) The influence of a relative movement among AAA ring and MTBD. Simulations with either a variable stalk-microtubule angle (left, grey) or with a fixed stalk-microtubule angle (left, orange) were run. (2) The influence of a distance-dependent bias to close the gap between the motor domains along the off-axis. Simulations with either a bias to step towards each other (right, grey) or without a bias to step towards each other (right, blue) were run. Note that the distance distribution for the flexible angle is from the same data as in b. The influence of other rules on dynein motility are shown in **Supplementary Figures 8**. (**b, c**) 100 simulations for each condition with more than 10,000 steps were performed. More details on the Monte Carlo simulation can be found in Materials and Methods, **Supplementary Figures 7, 8** and **Supplementary Movies 3-10**.

When we applied all these rules during the simulation, we could reproduce the experimental data for dynein stepping very well; the step size distributions of AAA rings and MTBDs of the simulation were almost identical to those observed in experimental data (**Supplementary Fig. 7**). Interestingly, certain parameters that were not provided in the model agreed well with previous experimental observations. For instance, we did not provide input regarding the spacing of the AAA rings, except for the experimentally determined step size distributions of the associated MTBDs and the relative movement (changes in angle ω) between AAA ring and MTBD (**Fig. 6 a**). Nevertheless, our simulation yielded inter-AAA ring distances (**Fig. 6 b**) that were very similar to those observed in early reported stepping experiments in which both AAA rings were labeled^29^. In addition, we also found good agreement for the probability of passing and not passing steps for the AAA rings when we compared it to prior experimental data^29^ without directly encoding this motion in the simulation (**Fig. 6 b**).

However, if we ignored any of the above listed rules during a Monte Carlo simulation, the simulated dynein motility did not match current or previous experimental observations (**Supplementary Fig. 8**). For instance, if we did not apply the tendency for the motor domains to step closer towards each other along the off-axis (**Fig. 6 c, Supplementary Fig. 8 b**), but rather allowed the motor domains to move in either direction along the off-axis, the motor domains drifted apart >100 nm in some simulations. Moreover, if we fixed the angle ω between the AAA ring and MTBD, the simulation produced a much larger inter-AAA ring distance (~27 nm) than experimentally measured (~18 nm)^29^ (**Fig. 6 c, Supplementary Fig. 6 d-l, Supplementary Movie 3, 9, 10**). Thus, encoding the experimentally derived set of rules and transition probabilities listed above is sufficient and necessary to recapitulate directed dynein motility using Monte Carlo simulations.

Since our three-color experimental data only captured the positions of one AAA ring and two MTBDs, the information regarding the location of the second AAA ring is missing. However, since our Monte Carlo simulation reproduced our and experimental data of others very well, we used this simulation to predict the positions of both AAA rings and MTBDs during motility. As a general validation of this approach, we compared the frequency of the experimental three-color domain orders (**Fig. 5**) to the frequency of three-color domain orders from the Monte Carlo simulations and found good agreement (**Supplementary Fig. 6 c**). Taking into account the four moving parts (2 AAA rings and 2 MTBDs in the homodimer), dynein can adopt 12 potential conformations (**Supplementary Fig. 6 a, b**). Of these 12 possible conformations, our simulation predicts that the first three conformations make up ~55% of the total, while the six least common confirmations comprised <20% (**Fig. 7, Supplementary Fig. 6 b**). Overall, conformations in which the stalks did not cross were more common (~76%) than conformations in which the stalks cross one another (~24%). Interestingly, the 8 conformations in which at least one motor domain has a stalk-microtubule angle of >90° were not predicted by previous models of dynein motility^1^ but make up ~65% of all dynein conformations based on our simulation. Taken together, our experimental data in combination with the Monte Carlo simulation provides a model of the distribution of dynein conformations during motility.

**Figure 7.**
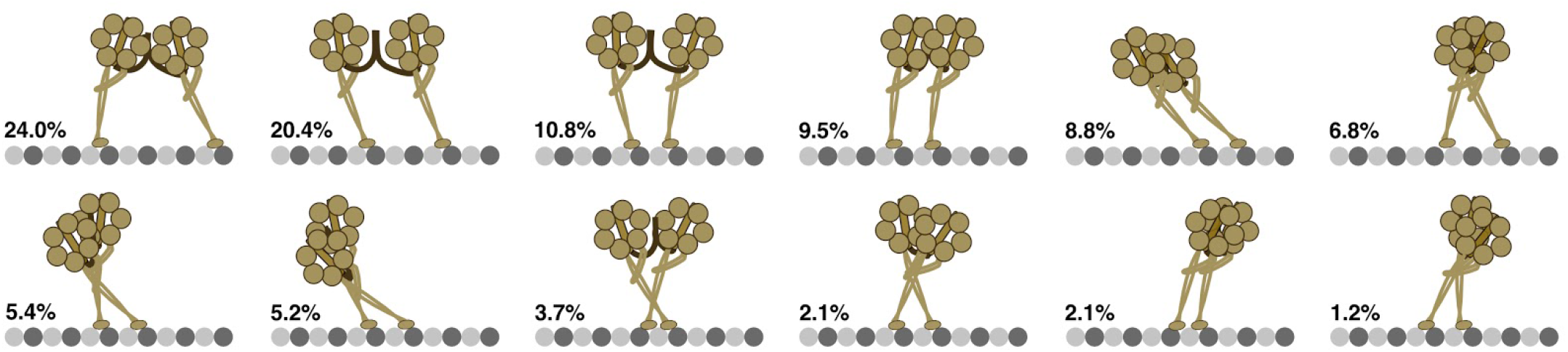
Model of Dynein Conformations. Monte Carlo simulation of dynein motility was used to determine the frequency of all 12 possible dynein conformational states (positions of AAA rings and MTBDs relative to the microtubule axis). Here, the occurrence of all 12 states is ordered from most common to least common (from top left to bottom right). Note that the absolute distance between the four domains (both AAA rings and both MTBDs) is irrelevant and that only the relative proximity of all four domains to the microtubule minus end determines the classification into conformational states. Moreover, we only show conformations for changes along the on-axis and are ignoring differences along the off-axis. A more detailed analysis of conformational states is given in **Supplementary Figure 6**.

## Discussion

Previous dynein stepping experiments have measured the positions of the AAA rings^29,30^. Here, we have been able to track the movements of both MTBDs, in combination with one AAA ring for the first time. Several technical challenges had to be overcome to make this measurement. First, the small 14 kDa MTBD^22^ had to be labeled without perturbing motor function. For this, the common HALO-tag^40^ and SNAP-tag^39^ are non-ideal, since they are twice as large as the MTBD. However, we found that a 14 amino acid-long YBBR-tag^41^ could be inserted into loop 5 of the MTBD and then labeled with a DNA FluoroCube^35^ without perturbing wild-type function. Second, tracking three colors for a prolonged time is not easily achievable with conventional dyes due to photobleaching, as even the photobleaching of one dye terminates the measurement. However, using DNA FluoroCubes, which are up to 50-fold more photostable than conventional organic dyes, enabled long-term tracking of many steps. Third, the distances between three colors had to be measured with nanometer accuracy. For this, we extended our previously published two-color data collection and imaging analysis pipeline^42^ to three-colors. All three technical advances described above were essential to accurately measure the positions of three domains of dynein as the motor undergoes hundreds of steps along a microtubule. These measurements provided new insights into the stepping behavior and conformational states of dynein, as discussed below.

### Flexibility within the motor domain allows dynein to adopt many conformational states

Our experimental data of the three-color-labeled dynein, combined with Monte Carlo simulations, show that dynein can adopt a large variety of conformations, many of which were not considered in prior stepping models of dynein^1,29,30^. These conformational states presented in **Figure 7** are most likely enabled by angular changes to the stalk that spans between the AAA ring and the MTBD. A wide range of stalk angles have been previously measured by electron microscopy^2,34^ and polarization microscopy^33^. However, the average angles between stalk and microtubule determined by cryo-electron microscopy by Imai et al.^34^ and Can et al.^2^ were ~42° and ~55°, respectively, and are smaller than what we measured for the leading (72°) and trailing (90°) motor domain. One explanation for this difference might be that our C-terminal fluorescent label on the AAA ring is not in the center but rather on the side of the AAA ring, which is closer towards the minus end and thus could bias the angle towards larger values. However, it is also possible that a moving dynein dimer adopts slightly different conformations than in rigor states, especially for monomers examined by electron microscopy.

In addition to measuring an average angle, we could obtain information on how the stalk angle is dependent upon the separation of the two MTBDs and other parameters that change as dynein steps (**Fig. 4, Supplementary Fig. 4, 5**). For instance, when the MTBDs are relatively close to each other (8 nm apart, the spacing between tubulin dimers), we found that the stalk-microtubule angles of both motor domains were relatively similar (**Supplementary Fig. 5 j-l**). However, if the MTBDs are further apart (>16 nm, two or more tubulin dimers), the motor domains tilt towards each other, resulting in a split-like conformation in which the AAA rings are closer together than the MTBDs (see **Figure 7** top most left and top most right state). Since we were able to track individual dyneins during many steps, we could also observe that the stalk-microtubule angle changes as dynein moves along its track. For example, when the trailing motor domain passed the leading motor domain, it often changed its angle from a steep to a more shallow angle (pivot-like motion) (**Fig. 3 c, d**). In summary, while other studies looked at distributions of static dynein and also observed flexibility within dyneins motor domain, we could for the first time observe the flexibility of an actively moving dynein, enabling us to gain new insights into motility.

By tracking one AAA ring and two MTBDs for the first time, we could derive more information on dynein conformational states during motility than could be inferred in prior studies that measured the AAA rings^29,30,32^. By combining our experimental data with Monte Carlo simulations, we could estimate the frequencies of 12 possible conformational relationships of the two AAA rings and the two MTBDs. Overall, this data suggests that states in which the MTBDs are further apart than the AAA rings are more common than vice versa. In addition, we find that for the leading motor domain, the MTBD is typically leading the AAA ring, while the trailing motor domain has an almost equal distribution of either AAA ring leading or MTBD leading. Moreover, our data show that the stalks of the dynein homodimer are rarely crossed; in other words, if the MTBD of motor domain 1 is leading the MTBD of motor domain 2, then it is very likely that the AAA ring of motor domain 1 is also leading the AAA ring of motor domain 2. Interestingly, we found that dynein only changes its conformational state after a step occurred in ~50% of all cases (**Fig. 4**), and that if dynein transitions between two states, it most likely switches between the two most common conformational states by a pivot-like transition (AAA ring and MTBD switch the lead) within the motor domain (**Fig. 5; Fig. 7**, two most upper left states). We could also demonstrate through simulation that the ability to adopt these conformational states requires a flexibility for the angle between the stalk and microtubule; if we fixed the angle between stalk and microtubule, as indicated in some dynein stepping models^1^, dynein can only adopt two of the twelve possible conformations (**Supplementary Fig. 6, Supplementary Movie 3, 9, 10**). Thus, our combination of three-color imaging and Monte Carlo simulation of dynein motility provides the first insights into the frequency and transitions between dynein conformational states.

### The AAA ring follows exploratory stepping of the MTBD

The stepping of the MTBD is initiated by ATP binding and a rotation of the AAA ring caused by the bending of the linker, dynein’s mechanical element^1,23,25–27^. The MTBD then is believed to execute a Brownian search followed by a rebinding to a new tubulin subunit. By tracking the MTBD and AAA ring simultaneously, we found that this search can have many different outcomes for the transition of the entire motor domain. In some cases, the MTBD can move (forward or backward) without a translation of the AAA ring and even if both domains step simultaneously, they can take differently sized steps (**Fig. 2, 3**).

These results are most consistent with flexibility within the dynein motor domain as opposed to rigid body movements of the motor domains. For example, relative movement between AAA ring and MTBD would allow the MTBD to take a short step, either backwards or forwards, while the AAA ring rotates and does not translate significantly along the long-axis direction (**Fig. 3 a**). However, if the MTBD takes a larger step after a Brownian search, the AAA ring on the same motor domain will be forced to follow because the step cannot be accommodated solely by an angular change between the AAA ring and MTBD. Previous studies suggested that only the AAA ring determines the step of the motor domain of dynein^29,30^. However, our data suggest that the AAA ring is essential to initiate and power the step, while the Brownian search of the MTBD and the flexibility of the stalk determines the step and which parts of the motor domain translocate (only MTBD or MTBD and AAA ring).

The sometimes differently sized steps of the AAA ring and the MTBD and the large variety of conformational states are strikingly different from other motor proteins such as kinesins, where the motor domain takes regular, 16-nm and almost exclusively forward steps^28,43^. Dynein’s inherent flexibility and ability to step in so many different ways might explain why a single dynein is more efficient than a single kinesin in circumventing obstacles such as microtubule associated proteins (MAPs)^44,45^.

### Dynamic, multi-color imaging is suitable for studying large molecular machines

Distance measurement between protein domains has been extensively investigated using Foerster resonance energy transfer (FRET)^46^. However, FRET is typically limited to short distances of 2-8 nm. In contrast, the multi-color measurements described here are not limited to any distance and thus can be applied to macromolecular complexes of any size to obtain direct distance relationships. Together with DNA FluoroCubes^35^, which provide a mechanism for following dynamics over long periods of time, the approach described in this work can be useful to investigate conformational changes of other multi-domain proteins or macromolecular complexes. For instance, the multi-color approach could be applied to study conformational changes of chaperones during the refolding of client substrates. We also envision that our approach could be applied to molecular machines that operate on other tracks than microtubules such as DNA. For example, the high-resolution multi-color approach could be used to investigate how chromatin remodelers interact with nucleosomes along DNA. Lastly, we anticipate that the framework provided in this work can be extended to four colors, which will further expand the reference points for investigating intra- and inter-protein dynamics.

## Acknowledgements

We are grateful to Christina Gladkova, Iris Grossman-Haham, and Zhen Chen (all University of California, San Francisco) for critical discussions of the manuscript. Andrew Carter (MRC Laboratory of Molecular Biology, UK) and Elizabeth Villa (University of California, San Diego) supplied the Matlab script for step detection of dynein. The authors gratefully acknowledge funding from the National Institutes of Health: R01GM097312, 1R35GM118106 (R.D.V., S.N.), and the Howard Hughes Medical Institute (N.S., N.Z., and R.D.V.).

## Author contributions

S.N. and R.D.V. designed the research; S.N. and N.Z. cloned, expressed and prepared samples; S.N. collected TIRF microscopy data; S.N. and N.S. developed data collection and analysis code; S.N. analyzed the data; S.N., N.S., and R.D.V. wrote the manuscript. All authors read and commented on the paper.

## Materials and Methods

### Flow-cell preparation

Flow-cells were assembled as previously described^36^. Briefly, we cut custom three-cell flow chambers out of double-sided adhesive sheets (Soles2dance, 9474-08×12 - 3M 9474LE 300LSE) using a laser cutter. We then used these three-cell flow chambers together with glass slides (Thermo Fisher Scientific, 12-550-123) and 170 μm thick coverslips (Zeiss, 474030-9000-000) to assemble the flow cells. Prior to assembly, coverslips were cleaned in a 5% v/v solution of Hellmanex III (Sigma, Z805939-1EA) at 50° C overnight and washed extensively with Milli-Q water afterwards.

### Assembly of DNA FluoroCubes for dynein labeling

FluoroCubes were assembled as previously described^35^. For each of the three six dye FluoroCubes, we used four 32 bp long oligonucleotide strands of which three were modified with two dyes and one with a functional tag; either a HALO-ligand^40^ for labeling of the HALO-tag or a Coenzyme A (CoA) for labeling of the YBBR-tag^41^ (**Supplementary Table 1**). The organic dye modified oligonucleotides were purchased from IDT and the oligonucleotides with functional tags were synthesized by Biomers. For each of the three FluoroCubes, four oligos were mixed to a final concentration of 10 μM each in folding buffer (5 mM Tris pH 8.5, 1 mM EDTA and 40 mM MgCl_2_). Then, we annealed the FluoroCubes by denaturation at 85° C for 5 min followed by cooling from 80° C to 65° C with a decrease of 1° C per 5 min. Afterwards the samples were further cooled from 65° C to 25° C with a decrease of 1° C per 20 min and finally held at 4° C. The folding products were purified and analyzed by 3.0% agarose gel electrophoresis in TBE (45 mM Tris-borate and 1 mM EDTA) with 12 mM MgCl_2_. The gel was run at 70 V for 2.5 hr on ice. Subsequently, we purified the DNA FluoroCubes by extraction and centrifugation in Freeze ’N Squeeze columns (BioRad Sciences, 732-6165). Prior to extraction, the gels were scanned using a Typhoon 9400 scanner (GE Healthcare).

### Dynein expression, purification, and labeling

We used recombinant *S. cerevisiae* cytoplasmic dynein (Dyn1) truncated at the N-terminus (1219-4093 aa) as a monomeric version expressed in a yeast strain with the following genotype MATa his3-11,5 ura3-1 leu2-3,112 ade2-1 trp-1 PEP4::HIS5 pGAL-ZZ-TEV-SNAPf-3XHA-D6-DYN1(MTBDL5:YbbR)-gsDHA for all our stepping experiments (VY1067^36^). This construct has a N-terminal SNAP-tag^39,41^, a C-terminal Halo-tag^40^ and a YBBR-tag^41^ inserted into loop 5 of the MTBD flanked by three glycines on either side (inserted as GGG-TVLDSLEFIASKLA-GGG between T3173 and L3174). In addition, we used VY208 (MATa his3-11,5 ura3-1 leu2-3,112 ade2-1 trp-1 PEP4::HIS5 pGAL-ZZ-TEV-sfGFP-3XHA-D6-DYN1-gsDHA) as a wild-type control for our velocity and processivity analysis. Expression of Dynein, either VY208 or VY1067, and yeast lysis were executed as previously described^15,36^.

For the VY1067 purification, the lysis supernatant was loaded onto an IgG Sepharose resin and washed with wash buffer (30 mM HEPES pH 7.4, 50 mM K-Acetate, 200 mM K-chloride, 2 mM Mg-Acetate, 1 mM EGTA pH 8.0, 1 mM ATP 10% Glycerol). Afterwards the resin was washed with TEV buffer (50 mM Tris-HCl pH 8.0, 150 mM K-Acetate, 6 mM Mg-Acetate, 1 mM EGTA pH 8.0, 1 mM ATP, 10% Glycerol). Then, we split the beads into two equal amounts to label each fraction (fraction A and B) differently. Fraction A was labeled with 5 μM of HALO ligand, ATTO 488 FluoroCubes and 5 μM CoA ATTO 647N FluoroCubes. Fraction B was labeled with 5 μM of CoA Cy3N FluoroCubes. For both labeling reactions, we added Mg-Acetate to a final concentration of 6 mM and EGTA pH 8.0 to a final concentration of 1 mM to the purified FluoroCubes perfore mixing it with the beads. Moreover, we added 2.5 μM of Sfp phosphopantetheinyl transferase to both reactions to enable YBBR-tag labeling^47^. Both reactions were incubated overnight. The next day, we washed both reactions extensively with TEV buffer. Then, we labeled fraction A and B with 20 μM of reverse complementary oligonucleotides in TEV Buffer for 12 hours. For fraction A we used the benzylguanine (BG) modified oligo: BG - GGT AGA GTG GTA AGT AGT GAA. And for fraction B we used the benzylguanine (BG) modified oligo: TTC ACT ACT TAC CAC TCT ACC - BG (both obtained from Biomers). Afterwards, both fractions were washed with additional TEV buffer. Next, we eluted the labeled proteins by incubating with 2 μM TEV protease in TEV buffer overnight. Finally, both samples were eluted and mixed to allow for dimerization.

Dynein with single organic dyes was prepared as FluoroCube labeled dynein, except that HALO Alexa 488 (Promega) was used instead of the HALO ATTO 488 FluoroCube, CoA 647 (NEB) was used instead of the CoA ATTO 647N FluoroCube, and CoA 547 (NEB) was used instead of the CoA Cy3N FluoroCube.

For the VY208 purification, the lysis supernatant was loaded onto a IgG Sepharose resin and washed with wash buffer. Afterwards, the resin was washed with TEV buffer. Then, dynein was eluted by incubating with 2 μM TEV protease in TEV buffer overnight.

### Microtubule preparation

The tubulin used in this work was purified as previously described^48^ We used unlabeled tubulin and biotinylated tubulin that were mixed at an approximate ratio of 20:1 in BRB80 (80 mM Pipes (pH 6.8), 1 mM EGTA, and 1 mM MgCl_2_). To start the polymerization reaction GTP was added to 1 mM and the solution was incubated for 15 min in a 37°C water bath. Then, 20 μM of Taxol (Sigma, T1912) was added and the mixture was incubated for another 2 hours at 37°C. At the start of each experiment, microtubules were spun over a 25% sucrose cushion in BRB80 at ~160,000 g for 10 min to remove unpolymerized tubulin and small filaments.

### Preparation of flow-cells with dynein

The flow chambers for the single-molecule assay were prepared as previously described^49^. To conduct all experiments described in this study, we prepared four slightly different types of environments: (1) For the majority of experiments we used biotinylated microtubules as tracks and a low ATP concentration (3 μM), (2) for one experiment we used axonemes as tracks and a low ATP concentration (3 μM) (**Supplementary Fig. 3**), (3) for another experiment we used biotinylated microtubules as tracks and added 1 mM ADP to rigorly bind dynein to microtubules (**Supplementary Fig. 2**), and (4) for one experiment we used biotinylated microtubules as tracks and a high ATP concentration (1 mM) (**Supplementary Fig. 2**).

For all microtubule-based experiments, we added 10 μl of 5 mg/ml Biotin-BSA in BRB80 and incubated for 2 min. Then, we washed with 20 μl of DAB (50 mM K-Ac, 30 mM HEPES, pH 7.4, 6 mM Mg(Ac)_2_, 1 mM EGTA) with 0.4 mg/ml κ-casein (Sigma, C0406). Next, we added 10 μl of 0.5 mg/ml Streptavidin in PBS and incubated for 2 min. Afterwards, we washed with 20 μl of DAB with 0.4 mg/ml κ-casein. Then, we added 10 μl of polymerized microtubules and incubated for 5 min. This was followed by a wash with 30 μl of DAB, 0.4 mg/ml κ-casein, and 10 μM Taxol.

For the axoneme-based experiment, we added 10 μl of unlabeled axonemes in BRB80 (80 mM Pipes (pH 6.8), 1 mM MgCl_2_, 1 mM EGTA) and incubated for 5 min. Then, we washed with 60 μl of DAB with 0.4 mg/ml κ-casein.

Once the tracks were added to the chambers, we could add dynein. Again, we added a few different dynein constructs: (1) For the majority of experiments we used dynein labeled with three-differently-colored DNA FluoroCubes, (2) for one experiment we used dynein labeled with three-differently-colored single dyes (**Supplementary Fig. 2**), and (3) for one experiment we used GFP-tagged dynein (**Supplementary Fig. 2**).

We added dynein diluted in DAB with 0.4 mg/ml κ-casein and 10 μM Taxol and incubated for 3 min. Afterwards we washed with 10 μl of DAB, 0.4 mg/ml κ-casein, and 10 μM Taxol. For the experiments used to extract dynein’s steps, we then added 10 μl of DAB, 0.4 mg/ml κ-casein, 10 μM Taxol, 3 μM ATP, an ATP regeneration system (1 mM phosphoenolpyruvate (Sigma, 860077), ~0.01 U pyruvate kinase (Sigma, P0294), ~0.02 U lactate dehydrogenase (Sigma, P0294)), and the PCA/PCD/Trolox oxygen scavenging system^50,51^. For the velocity and processivity comparison of FluoroCube-labeled and GFP-tagged dynein (**Supplementary Fig. 2**), we added 10 μl of DAB, 0.4 mg/ml κ-casein, 10 μM Taxol, 1 mM ATP, and the PCA/PCD/Trolox oxygen scavenging system^50,51^. Lastly for the brightness and photostability comparison of FluoroCube-labeled and single dye-labeled dynein (**Supplementary Fig. 2**) we added 10 μl of DAB, 0.4 mg/ml κ-casein, 10 μM Taxol, 1 mM ADP, and the PCA/PCD/Trolox oxygen scavenging system^50,51^.

We note that using κ-casein was essential as other caseins such as β-casein made FluoroCube-labeled dynein stick to the glass and not move along microtubules or axonemes. Moreover, we note that the concentration of the PCA/PCD/Trolox oxygen scavenging system^50,51^ is very important to achieve optimal photostability. We used the following concentrations in all our experiments: 2.5 mM of protocatechuic acid (PCA) (Sigma: 37580) at pH 9.0, 5 units of protocatechuate-3,4-dioxygenase (PCD) (Oriental yeast company Americas Inc.: 46852004), and 1 mM Trolox (Sigma: 238813) at pH 9.5.

### Fluorescent beads for image registration

To register the three channels, we used TetraSpeck™ beads (Thermo Fisher Scientific, T7279) based on a previously described protocol for two-color image registration^36^. To this end, we prepared one of the three flow chambers of our flow cells with dynein (or with the DNA-origami nanoruler (**Supplementary Fig. 1**)). and another flow chamber on the same flow cell with TetraSpeck™. The beads were immobilized by adding 10 μl of 1 mg/ml Poly-D-lysine (Sigma, P6407) in Milli-Q water to the flow-cell, followed by a 3 min incubation and a wash with 20 μl of BRB80 (80 mM Pipes (pH 6.8), 1 mM EGTA, and 1 mM MgCl_2_). Afterwards, we added 10 μl of 1:300 diluted TetraSpeck™ beads in BRB80 and incubated for 5 min. Finally, the flow-cell was washed with 40 μl of BRB80.

### DNA-origami nanoruler distance measurements

We designed and assembled DNA-origami nanorulers based on a previously described protocol^52^. This particular three-color nanoruler design is based on the 12 helix bundle (12 DNA double helices) and is assembled with fluorescently labeled oligos with one dye of each ATTO 488, Cy3, and ATTO647N per ruler. Moreover, biotinylated oligos are incorporated into the structure on the opposite side of the fluorescent dyes to enable surface immobilization (**Supplementary Fig. 1**, **Supplementary Table 2**). We designed the ruler in such a way that the ATTO 488 and the ATTO 647N dye are separated by ~28 nm, the ATTO 488 and the Cy3 dye are separated by ~14 nm, and the Cy3 and the ATTO 647N dye are separated by ~14 nm.

For assembly of three-color nanorulers, oligos were mixed to a final concentration of 200 nM each in folding buffer (5 mM Tris pH 8.5, 1 mM EDTA and 22 mM MgCl_2_) with 20 nM of p8064 scaffold (Tilibit; Single-stranded scaffold DNA, type p8064). Then, nanorulers were annealed by first denaturing at 85° C for 5 min followed by cooling from 80° C to 65° C with a decrease of 1° C per 5 min. Afterwards the samples were further cooled from 65° C to 25° C with a decrease of 1° C per 30 min and finally held at 4° C.

Three-color nanorulers were purified by agarose gel electrophoresis. Structures were loaded into 2% agarose gels and run at 70 V for ~2 hrs in TBE buffer (45 mM Tris, 45 mM boric acid, 1 mM EDTA) supplemented with 11 mM MgCl2. Bands corresponding to well folded monomeric three-color nanorulers were excised, crushed and spun through a Freeze N’ Squeeze column (BioRad Sciences, 732-6165) for 3 min at 13,000g at 4°C. Prior to extraction, the gels were scanned using a Typhoon 9400 scanner (GE Healthcare). The three-color DNA-origami nanorulers were stored at 4°C.

Flow-cells with nanorulers were prepared as follows: we first added 10 μl of 5 mg/ml Biotin-BSA in BRB80 and incubated for 2 min. Then, we washed with 20 μl of PBS (pH 7.4), added 10 μl of 0.5 mg/ml Streptavidin in PBS (pH 7.4) and incubated for another 2 min. Afterwards, we washed with 20 μl of PBS (pH 7.4) supplemented with 10 mM MgCl_2_. Then, we added 10 μl of three-color nanoruler and incubated for 5 min. Finally, we washed with 30 μl of PBS (pH 7.4) supplemented with 10 mM MgCl_2_ and then added the PCA/PCD/Trolox oxygen scavenging system^50^ in PBS (pH 7.4) supplemented with 10 mM MgCl_2_.

### Microscope setup

Data collections for all experiments were carried out at room temperature (~23° C). For imaging, we used a total internal reflection fluorescence (TIRF) inverted microscope (Nikon Eclipse Ti microscope) equipped with a 100× (1.45 NA) oil objective (Nikon, Plan Apo λ). Moreover, we used two Andor iXon 512×512 pixel EM cameras, DU-897E and no additional magnification, resulting in a pixel size in the image plane of 159 nm. The microscope is also equipped with two stepping motor actuators (Sigma Koki, SGSP-25ACTR-B0) mounted on a KS stage (KS, Model KS-N) and a custom-built cover to reduce noise from air and temperature fluctuations. In addition, a reflection based autofocus unit (FocusStat4) was custom adapted to our TIRF microscope (Focal Point Inc.). We used a 488 nm laser (Coherent Sapphire 488 LP, 150 mW), a 561 nm laser (Coherent Sapphire 561 LP, 150 mW), and a 640 nm laser (Coherent CUBE 640-100C, 100 mW) for data collection. The laser lines combined, passed through an AOTF to control illumination power, were 6-fold enlarged, passed through a quarter wave plate (ThorLabs, AQWP05M-600) and then focused using an achromatic doublet f=100 mm (Thorlabs) on a conjugate back focal plane of the objective outside of the microscope. We adjusted the TIRF angle by moving a mirror and focusing lens simultaneously. In the upper turret of the microscope, we mounted a TIRF cube containing excitation filter (Chroma, zet405/491/561/638x), dichroic mirror (zt405/488/561/638rpc), and emission filter (Chroma, zet405/491/561/647m). The lower turret contained a filter cube (Chroma, TE/Ti2000_Mounted, zet 405/488, T560lpxr, zet 561/640m) that directs ATTO 488 emission towards the back camera and the Cy3N as well as the ATTO 647N emission towards the left camera. The acquisition software was μManager^53^ 2.0.

### Single-molecule TIRF data collection

All TetraSpeck™ bead, nanoruler, and dynein datasets were acquired with a ‘16 bit, conventional, 3 MHz’ camera setting, a preamp gain of 5x in conventional CCD mode (i.e., no EM gain), using overlapped mode (i.e. exposure of the next image starts while the readout of the CCD takes place). The intensity (irradiance) at the objective was 120 W/cm^2^ (488 nm laser), 120 W/cm^2^ (561 nm laser), and 160 W/cm^2^ (640 nm laser).

The exposure TTL signal of the camera on the left was connected to the input trigger TTL connector of the camera on the back, as well as to the input connector of a microcontroller used to control laser light intensity through the AOTF. As a result of this configuration, when the camera on the left runs a sequence in internal trigger mode, the camera in the back is ready to be triggered only every other image (i.e. it will expose during each odd numbered exposure of the camera on the left).

For data collection of dynein stepping, we prepared one chamber with TetraSpeck™ beads and another chamber on the same microscopy slide with three-color dynein either moving along microtubules or axonemes. Every data collection cycle was started by imaging a 20 by 20 grid of TetraSpeck™ beads. To do so, we collected micrographs with alternating excitation with the 488 nm, 561 nm, and 640nm laser (110 msec exposure each) at positions about 1 micron apart. After imaging at one position, we moved the stage and waited 3 sec before collecting data at the new position to minimize drift effects.

Once the TetraSpeck™ beads dataset was collected, we moved to the chamber with the three-color dynein and acquired six, 500-frame-long movies with 110 msec exposure times using a Beanshell script for Micro-Manager^53^, that resulted in repeating sequences of exposure with the 488 nm laser collected from the back camera, 561 nm laser collected on the left, 640 nm laser collected on the left, followed by a necessary dummy image (which was discarded), 561 nm laser collected on the left, and 640 nm laser collected also on the left (**Supplementary Fig. 3**). Thus, we collected one image every 110 msec on one of the two cameras connected to our TIRF microscope. Since we wanted to read out data continuously and one camera triggered the readout of the other, we reached a point every other cycle in which both cameras were busy and not able to record new images. Even though this resulted in one blank image every six images it was far better than having the cameras trigger via the software as this resulted in much longer dead time between frames and effective interval times of ~200 msec per image compared to the 110 msec per image that we were able to achieve with this setup. Together, this led to the following: the blue and red channel were acquired every cycle which resulted in a 330 msec interval, while the green channel had to be skipped every other cycle leading to an interval of 660 msec. After collecting the dynein movies, we moved back to the TetraSpeck™ beads chamber to collect another 20 x 20 grid, which was used as a control to test whether any changes in image registration occurred during acquisition (see **Supplementary Fig. 1**). We only accepted datasets if σ_reg_ < 1 nm.

For the single color stepping traces with continuous illumination (**Supplementary Fig. 3**), we did not acquire any TetraSpeck™ beads but only acquired dynein movies with 1,500 frames total, with an exposure time of 110 msec.

For the data collection of the DNA-origami nanorulers (**Supplementary Fig. 1**), we acquired 20 movies with an alternating exposure of 400 msec between all three channels. We also acquired images of TetraSpeck™ beads before and after the data collection of the DNA-origami nanorulers to perform image registration and to test the registration, respectively.

For the velocity and processivity comparison (**Supplementary Fig. 2**), we did not acquire any TetraSpeck™ beads but only acquired dynein movies with 100 frames total, with an exposure time of 110 msec and a 5 sec interval between acquisition sequences.

For the brightness and photostability comparison (**Supplementary Fig. 2**), we did not acquire any TetraSpeck™ beads but only acquired dynein movies with 200 frames total with an exposure time of 110 msec and a 3.1 sec interval between acquisition sequences.

### Velocity and processivity analysis

Data for three-color-labeled dynein and GFP-tagged wild-type dynein (**Supplementary Fig. 2**) was acquired using μManager^53^ 2.0. Subsequently, the data was analyzed in ImageJ^54^ by generating kymographs and then measuring displacement as a function of time.

### Bleaching analysis of three-color dynein

Data for three-color dynein labeled with either FluoroCubes^35^ or conventional single dyes (**Supplementary Fig. 2**) was acquired using μManager^53^ 2.0. Subsequently, single molecules were localized using the Spot Intensity Analysis plugin in ImageJ^54^ with the following settings: Time interval of 3.1 sec, Electron per ADU of 1.84, Spot radius of 3, Noise tolerance of 100 for the FluoroCube data and of 50 for the conventional dye data, and a Median background estimation. The number of frames to check was set to 20 for the FluoroCube data and 10 for the conventional dye data. Afterwards the data was plotted using a custom python script as previously described^35^.

### Single-molecule TIRF data analysis of dynein stepping

For the three-color dynein stepping analysis, the emitters of dynein and TetraSpeck™ beads were fitted and localized using the μManager^53^ “Localization Microscopy’ plug-in (**Supplementary Table 3**). After localizing all probes, we registered the three channels using the same affine-based approach as previously described for two colors^36^ (**Supplementary Fig. 1**). Then, tracks of individual motors were extracted using the μManagers^53^ “Localization Microscopy’ plug-in. To this end, we set the minimum frame number to 125, the maximum number of missing frames to 350, the maximum distance between frames to 200 nm and the total minimum distances of the full track to 200 nm. If one trace of a three-color trace had a frame, which was not detected (either because of blinking or because the photon output was below our threshold for detection) we decided to keep these traces but ignore the particular frame in our stepping analysis. Afterwards, we rotated tracks of individual motors using a principal component analysis (PCA) implemented in python. Next, we applied a custom Matlab (Matlab R2019b) script to identify individual steps using Chung-Kennedy edge-detecting algorithm with a window size of 2 as previously described^29^ and further analyzed the data in a custom written python script. Only steps for which the step itself and the previous as well as the following step had a standard error of the mean of the 2D distance of less than 8 nm were considered for further analysis. Moreover, we only called steps if a color moved >4 nm along either on- or off-axis. Lastly, once the steps were detected, we manually inspected each stepping trace to identify anomalies. This was very important as the step detection algorithm sometimes detected more steps that we would have identified manually. The main reason for this is because the individual motors move at various velocities and thus, some motors take more steps during a time window than others. However, since we used the same window size for step detection, the slow motors are more likely to have additional steps detected by the algorithm.

For the single color acquisition as shown in **Supplementary Figure 3**, we also extracted spots using the μManagers^53^ “Localization Microscopy’ plug-in but did not perform any image registration. To extract single traces we set the minimum frame number to 750, the maximum number of missing frames to 750, the maximum distance between frames to 200 nm and the total minimum distances of the full track to 500 nm. Afterwards we analyzed data as described for the three-color data except that we only accepted steps if one color moved >5 nm and that we used a window size of 6 rather than 2 (frequency of imaging the same color during an acquisition cycle was three times faster than for the three color acquisition). For the three-color data shown in **Supplementary Figure 3** (which is the same as used throughout the manuscript) we performed the exact same analysis except that we used a window size of 2 since the effective imaging interval was three times shorter than for the single color data acquisition. For the single color acquisition as well as the axonemal acquisition in comparison to the microtubule acquisition, we decided to use the fully automated detection and no manual correction of these stepping traces to avoid any bias.

To compare axonemal and microtubule stepping data (**Supplementary Fig. 3**), we followed the same protocol as described for our three-color stepping analysis. However, we only accepted steps if one color moved >5 nm and decided to use the fully automated detection and no manual correction of these stepping traces to avoid any bias.

### Image registration and distance measurements for DNA-origami nanoruler

Image registration and distance measurements between multiple dyes on the DNA-origami nanoruler (**Supplementary Fig. 1**) were carried out as previously described^36^. Since this is a three-color dataset instead of a two-color dataset, we carried out the distance measurements for individual spot pairs (e.g. Cy3 and ATTO 488 or Cy3 and ATTO 647N or ATTO 488 and ATTO 647N). To localize individual spots (**Supplementary Table 3**) and to extract spots which contained a nanoruler with all three labels, we used μManagers^53^ “Localization Microscopy’ plug-in. To this end, we set the minimum frame number to 18, the maximum number of missing frames to 2, the maximum distance between frames to 15 nm, the total minimum distances of the full track to 0 nm, and the maximum distances between each dye pair to 90 nm.

### Negative stain electron microscopy data collection of three-color nanoruler

For negative-stain electron microscopy, agarose gel-purified nanorulers were incubated on freshly glow discharged carbon coated 400 mesh copper grids for 1 min. Afterwards the sample was blotted off and a 0.75% uranyl formate solution was applied immediately for staining and blotted off without incubation. This staining was repeated four times and followed by a last incubation for which the stain was incubated for 45 sec before blotting. Samples were air dried before imaging. The data were collected at UCSF, on a Tecnai T12 microscope operating at 120 kV, using a 4k×4k CCD camera (UltraScan 4000, Gatan).

### Monte Carlo simulation of dynein stepping

For the Monte Carlo simulation we used our experimental data as input. Briefly, we defined a start condition (position of both MTBDs), followed by a loop of simulations for continuous stepping which ended as soon as dynein reached the minus end of the microtubule lattice. The stepping loop was set up as follows: (1) Determine the relative position (left or right, leading or trailing) of both MTBDs and the distance between both MTBDs, (2) based on the inter-MTBD distance and the probability for leading and trailing MTBD to step, determine if leading or trailing MTBD is taking the next step, (3) based on the inter-MTBD distance, determine the dwell time for the MTBD that will take the next step, (4) based on the relative position of the MTBD that is taking the next step (left or right, leading or trailing) and based on the inter-MTBD distance, determine the step size, (5) based on the MTBD step size and the current and future (after the MTBD stepped) relative position of AAA ring and MTBD, determine the step size of the AAA ring. If the end of the microtubule is not reached, start over at step (1). The parameters such as length and number of protofilaments of the microtubule lattice can be predefined. For all our simulations we used a microtubule lattice with 13 protofilaments and a length of 79 tubulin dimers (~630 nm). Note that during the simulation the MTBDs are the main driver of motility and we let the AAA rings follow by a defined set of rules as described above. For the simulations in which we removed some of our predefined rules, we used slightly different data as input.

### Figure preparation

All figures and graphs were prepared by either using ImageJ (light microscopy data), Affinity designer (version 1.6.1, Serif (Europe) Ltd), or Python (version 2.7, Python Software Foundation).

### Statistics

We discussed the inherent uncertainty due to random or systematic errors for each result and their validation in the relevant sections of the manuscript. Moreover, we included details about sample size, number of independent calculations, and the calculation of the error bars in the figures or in the respective figure captions.

