## Supplementary Information for "Three-color single-molecule imaging reveals conformational dynamics of dynein undergoing motility"

*for*

This document includes:

Supplementary Information Figures (Supplementary Figure 1 - 9)

Supplementary Information Tables (Supplementary Table 1 - 3)

References for the Supplementary Information

### Supplementary Information Figures

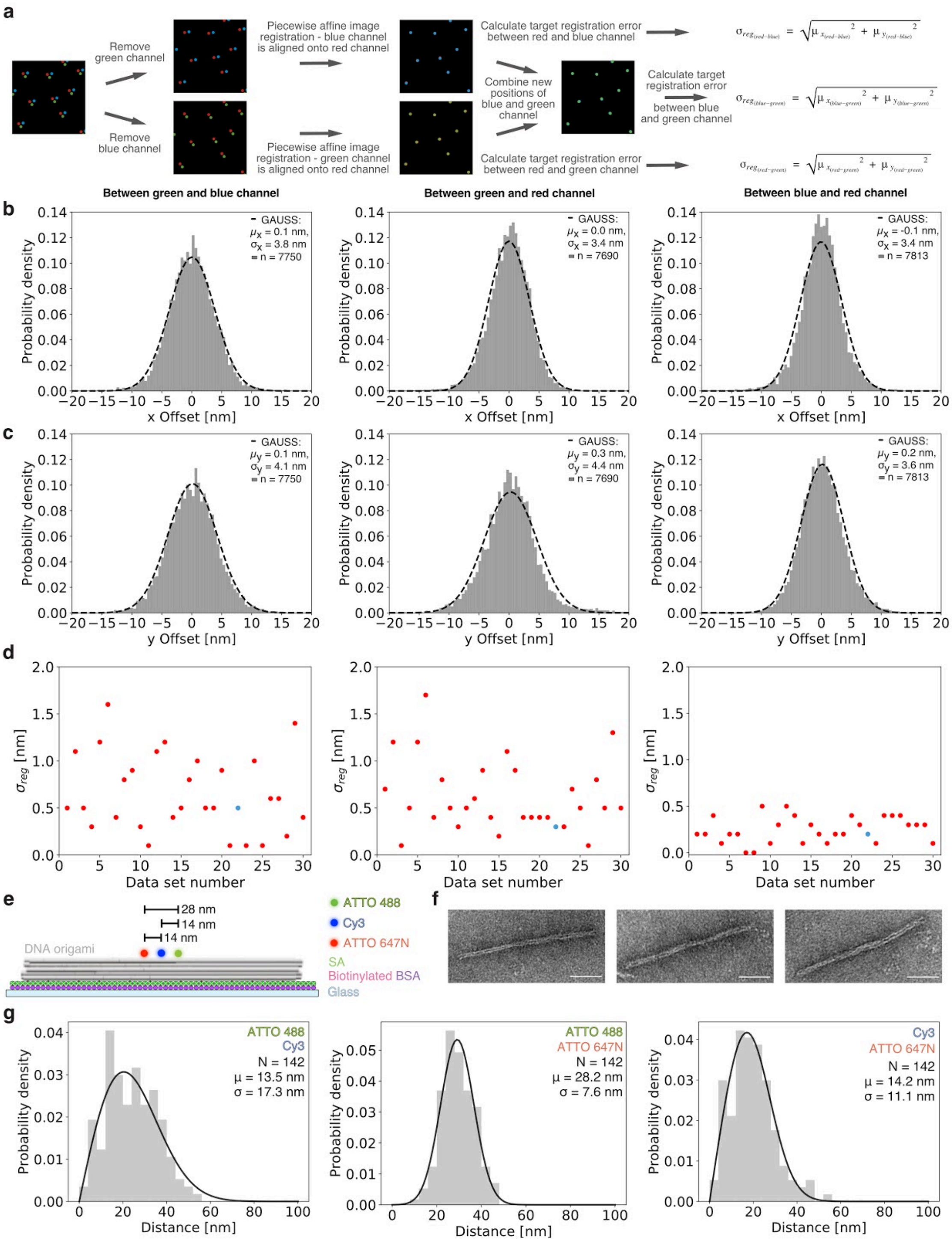


**Supplementary Figure 1 | Example and validation of three-color image registration.** (**a**) Workflow for three-color image registration. Tetraspeck beads are imaged in three channels (green, blue, and red) and images are divided into combinations of blue and red (top) or green and red channels (bottom). Then, both two-color datasets are aligned as previously described[1]. For both, the red channel serves as the reference and only blue and green positions are shifted. Afterwards the target registration error for all three color combinations is calculated. Note, to calculate the registration error between green and blue, the two split, registered data sets are combined and the red channel is removed. (**b-d**) Example results of image registration for all three color combinations are shown: green and blue channel (left panels), green and red channel (middle panels), and blue and red channel (right panels). Histogram of (**b**) x-axis and (**c**) y-axis offset after image registration with Gaussian fit (dashed black line). Sample size (n), average offset (μ) and its standard deviation (σ) are given. (**d**) Registration accuracy $\sigma_{reg}$ is shown for 30 datasets. The blue dot is the registration error of the data shown in detail in **b** and **c**. For the three-color dynein stepping experiments we accepted datasets if $\sigma_{reg} < 1$nm.

(**e-g**) Validation of three-color image registration with DNA origami nanorulers. (**e**) Schematic of the design of a three-color DNA-origami based nanoruler[2,3]. On this nanoruler, the three dyes ATTO 488, Cy3, and ATTO 647N are placed at well-defined distances. On the opposite side of the dyes, the nanoruler is functionalized with biotin for immobilization on a glass coverslip via biotinylated BSA and streptavidin (SA). (**f**) Negative stain electron microscopy micrographs of three-color nanoruler. Scale bar is 50 nm. (**g**) Histogram of measured distances for all three color combinations. The black line shows a fit with a probability distribution function described in[1] with the average distance (μ), its standard deviation (σ) and the sample size (N). Note, the standard deviation of the ATTO 488 and Cy3 color combination is higher than that of the other two color combinations because ATTO 488 and Cy3 yielded a larger localization error than color combinations that included the well-localized ATTO 647N dye. The data shown is for a single acquisition that was successfully reproduced five times (not shown).

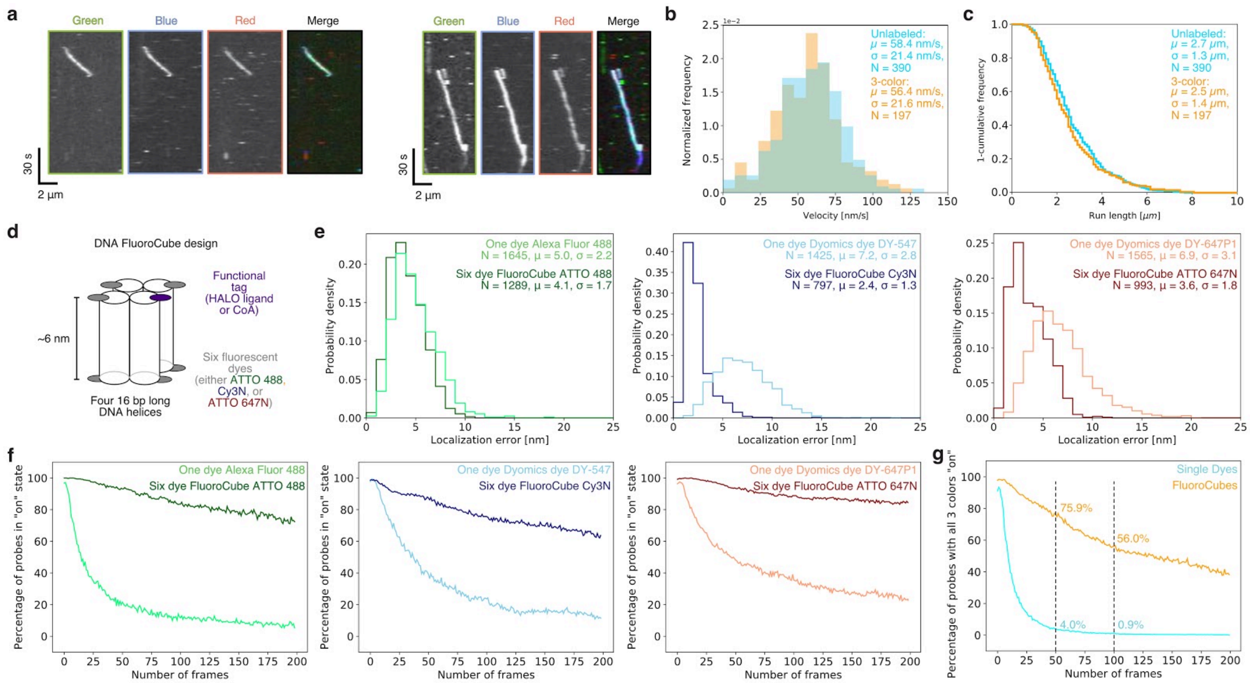

**Supplementary Figure 2 | Validation and benefits of FluoroCube labeled three-color dynein.** (**a-c**) Comparison of velocity and processivity of three-color dynein labeled with individual organic dyes or six dye FluoroCubes. (**a**) Example kymographs of three-color dynein. (**b**) Velocity histogram with average velocity ($\mu$) and its standard deviation ($\sigma$) for three-color dynein labeled with six dye DNA FluoroCubes[4] (orange) or GFP-tagged wild-type dynein (cyan). (**c**) A '1-cumulative frequency distribution plot' of run length with average length ($\mu$) and its standard deviation ($\sigma$) for three-color dynein labeled with six dye DNA FluoroCubes (orange) or GFP-tagged wild-type dynein (cyan). (**b, c**) The data is pooled from many movies with motors from the same preparation but with freshly prepared microscope slides (n=3) and the sample size N is given.

(**d-g**) Comparison of photostability and localization error of rigor-bound dynein labeled with organic dyes or six dye FluoroCubes. (**d**) Design of DNA FluoroCubes as previously described[4]. Here we used either a HALO ligand or CoA as functional tags and either six ATTO 488 dyes, six Cy3N dyes, or six ATTO 647N dyes. (**e-g**) Labeling of three-color dynein heterodimers with FluoroCubes[4] and dyes was as follows: For one motor domain the AAA ring is labeled via a HALO-tag[5] with a six dye ATTO 488 FluoroCube (dark green) or an Alexa Fluor 488 (light green) and the MTBD is labeled via YBBR-tag[6] with a six dye ATTO 647N FluoroCube (dark red) or a Dyomics dye DY-647P1 (light red). For the other motor domain only the MTBD is labeled via YBBR-tag[6] with a six dye Cy3N FluoroCube (blue) or a Dyomics dye DY-547 (light blue). For the measurement of photostability and localization error a three-color dynein is rigor-bound to microtubules using 1 mM ADP. Note that the photostability is likely slightly higher than measured as some dyneins may have detached prior to photobleaching. (**e**) Localization error for respective probes when attached to rigor-bound dynein. The data is pooled from many movies with motors from the same preparation but with freshly prepared microscope slides (n=3). Sample size (N), average localization error (μ) and its standard deviation (σ) are given. (**f**) Photostability of single-organic dyes compared to DNA FluoroCubes. Here, the number of probes in the on state is shown as a function of frames recorded. (**g**) Cumulative photostability of all three-color dyneins either labeled with single-organic dyes or with DNA FluoroCubes. Here, the number of dyneins for which all probes are in the on state is shown as a function of frames recorded. (**f, g**) Number of probes measured is given in **e**. Details about imaging conditions are given in **Materials and Methods**.

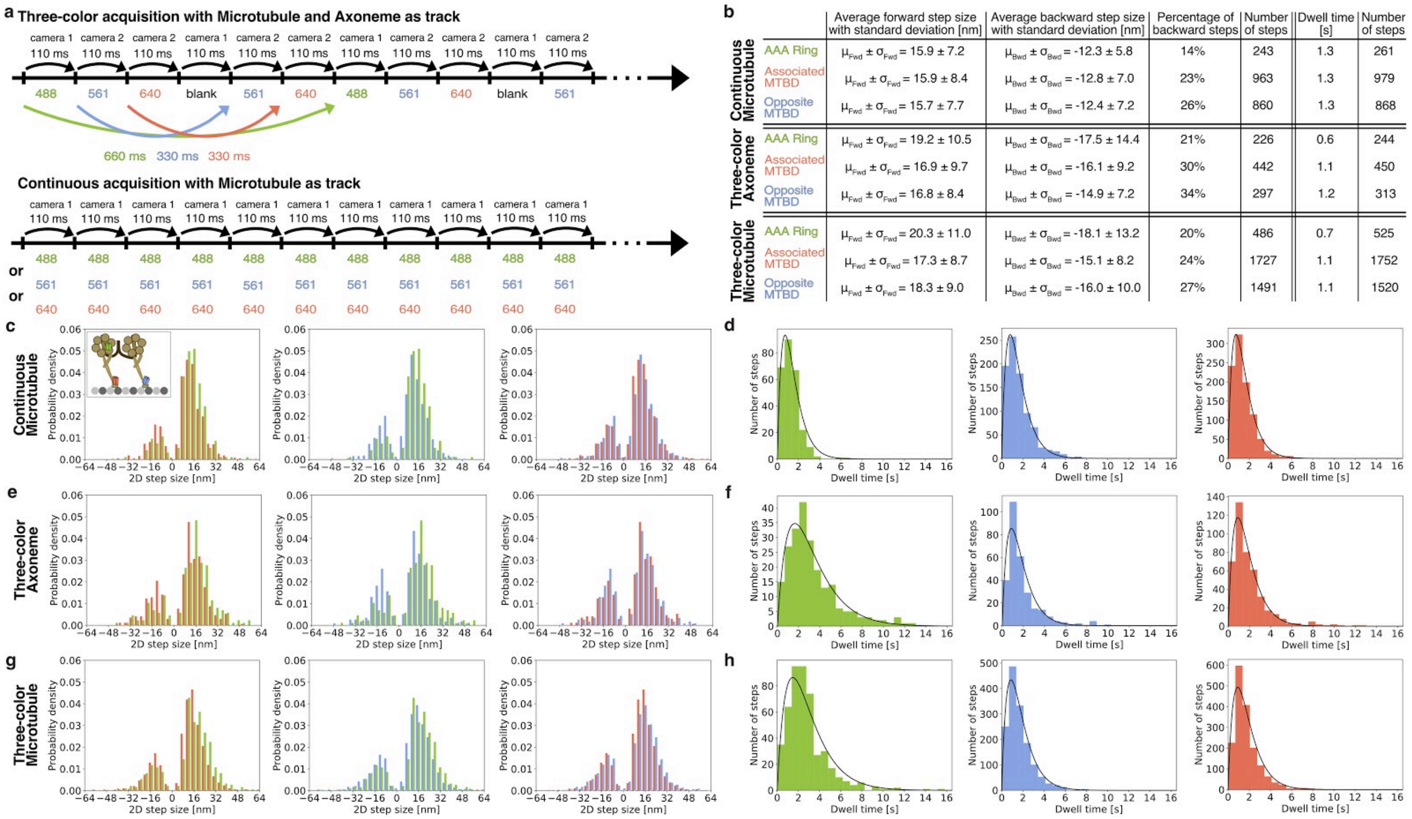


| | | Average forward step size with standard deviation [nm] | Average backward step size with standard deviation [nm] | Percentage of backward steps | Number of steps | Dwell time [s] | Number of steps |
|---|---|---|---|---|---|---|---|
| Continuous Microtubule | AAA Ring | $\mu_{Fwd} \pm \sigma_{Fwd} = 15.9 \pm 7.2$ | $\mu_{Bwd} \pm \sigma_{Bwd} = -12.3 \pm 5.8$ | 14% | 243 | 1.3 | 261 |
| | Associated MTBD | $\mu_{Fwd} \pm \sigma_{Fwd} = 15.9 \pm 8.4$ | $\mu_{Bwd} \pm \sigma_{Bwd} = -12.8 \pm 7.0$ | 23% | 963 | 1.3 | 979 |
| | Opposite MTBD | $\mu_{Fwd} \pm \sigma_{Fwd} = 15.7 \pm 7.7$ | $\mu_{Bwd} \pm \sigma_{Bwd} = -12.4 \pm 7.2$ | 26% | 860 | 1.3 | 868 |
| Three-color Axoneme | AAA Ring | $\mu_{Fwd} \pm \sigma_{Fwd} = 19.2 \pm 10.5$ | $\mu_{Bwd} \pm \sigma_{Bwd} = -17.5 \pm 14.4$ | 21% | 226 | 0.6 | 244 |
| | Associated MTBD | $\mu_{Fwd} \pm \sigma_{Fwd} = 16.9 \pm 9.7$ | $\mu_{Bwd} \pm \sigma_{Bwd} = -16.1 \pm 9.2$ | 30% | 442 | 1.1 | 450 |
| | Opposite MTBD | $\mu_{Fwd} \pm \sigma_{Fwd} = 16.8 \pm 8.4$ | $\mu_{Bwd} \pm \sigma_{Bwd} = -14.9 \pm 7.2$ | 34% | 297 | 1.2 | 313 |
| Three-color Microtubule | AAA Ring | $\mu_{Fwd} \pm \sigma_{Fwd} = 20.3 \pm 11.0$ | $\mu_{Bwd} \pm \sigma_{Bwd} = -18.1 \pm 13.2$ | 20% | 486 | 0.7 | 525 |
| | Associated MTBD | $\mu_{Fwd} \pm \sigma_{Fwd} = 17.3 \pm 8.7$ | $\mu_{Bwd} \pm \sigma_{Bwd} = -15.1 \pm 8.2$ | 24% | 1727 | 1.1 | 1752 |
| | Opposite MTBD | $\mu_{Fwd} \pm \sigma_{Fwd} = 18.3 \pm 9.0$ | $\mu_{Bwd} \pm \sigma_{Bwd} = -16.0 \pm 10.0$ | 27% | 1491 | 1.1 | 1520 |

**Supplementary Figure 3 | Validation of three-color image acquisition approach.** (**a**) Top: Schematic illustration of data collection in three-color acquisition mode minimizing dead time between frames. The exposure signal of the camera on the back (camera 1) of the microscope triggered exposure of the camera on the left (camera 2); this arrangement resulted in the back camera exposing only every other image (i.e. every odd numbered exposure of the left camera coincided with an exposure of the back camera). To reduce background, it was essential to illuminate only one laser for each channel, and to maintain image registration, it was essential to not have any moving filters in the system. We used the illumination sequence depicted in the figure. This led to the following rates: the blue and red channels were acquired every cycle, which resulted in 330 ms intervals, while the green channel was skipped every other cycle leading to 660 ms intervals. Bottom: Acquisition cycle for data collection of a single color. In this mode movies for a single color were collected continuously from a single camera with a 110 ms exposure time and interval. (**b**) Table summarizing the average values of step size and dwell time data of three-color dynein either collected in continuous single-color mode (moving along microtubules) or in three-color acquisition mode (either on microtubules or on axonemes). (**c, e, g**) Step size distributions of green-labeled AAA ring, blue-labeled MTBD, and red-labeled MTBD either collected in (**c**) continuous single-color mode moving on microtubules, in three-color acquisition mode moving along (**e**) axonemes or (**g**) microtubules. Left: Histogram of on-axis step sizes of dynein's AAA ring (green) and the MTBD (red) of the same motor domain (associated MTBD). Middle: Histogram of on-axis step sizes of dynein's AAA ring (green) and the MTBD (blue) on the opposite motor domain. Right: Histogram of on-axis step sizes of dynein's MTBDs (blue and red). (**c**) Box in the top left shows a schematic of three-color dynein. (**d, f, h**) Histogram of the dwell times of green-labeled AAA ring, blue-labeled MTBD, and red-labeled MTBD either collected in the (**d**) continuous single-color mode moving on microtubules, in the three-color acquisition mode moving along (**f**) axonemes or (**h**) microtubules. The black lines are fits of a convolution of two exponential functions with equal decay constants.

For the data presented in this work we acquired images in all three channels in an alternating fashion (see panel **a**). However, to enable fast acquisition with minimal dead time, we skipped the green channel every other round. This data shows that the three-color acquisition sequence did not change the step size distributions when compared to single-color, continuous acquisition. For instance, for both modes we found that the MTBDs take more backward than forward steps than the AAA ring. However, the dwell time for the continuous acquisition of the AAA ring was the same as for the MTBDs while it was longer for the AAA ring compared to the MTBDs in the three-color acquisition.

Moreover, this data shows that stepping along microtubules and axonemes of the three-color dynein is very similar. The largest difference among these two datasets was observed for the percentage of backward steps of the MTBDs. Here, it was more likely that the MTBDs take backward steps when moving along axonemes (30% and 34%) compared to when moving along microtubules (24% and 27%).

We note that the stepping traces for this data were not manually inspected (and corrected if necessary), leading to a higher percentage of shorter forward and more backward steps because the step detection algorithm called more steps than we would have done manually. Nevertheless, we decided to use the fully automated detection without manual correction to avoid bias.

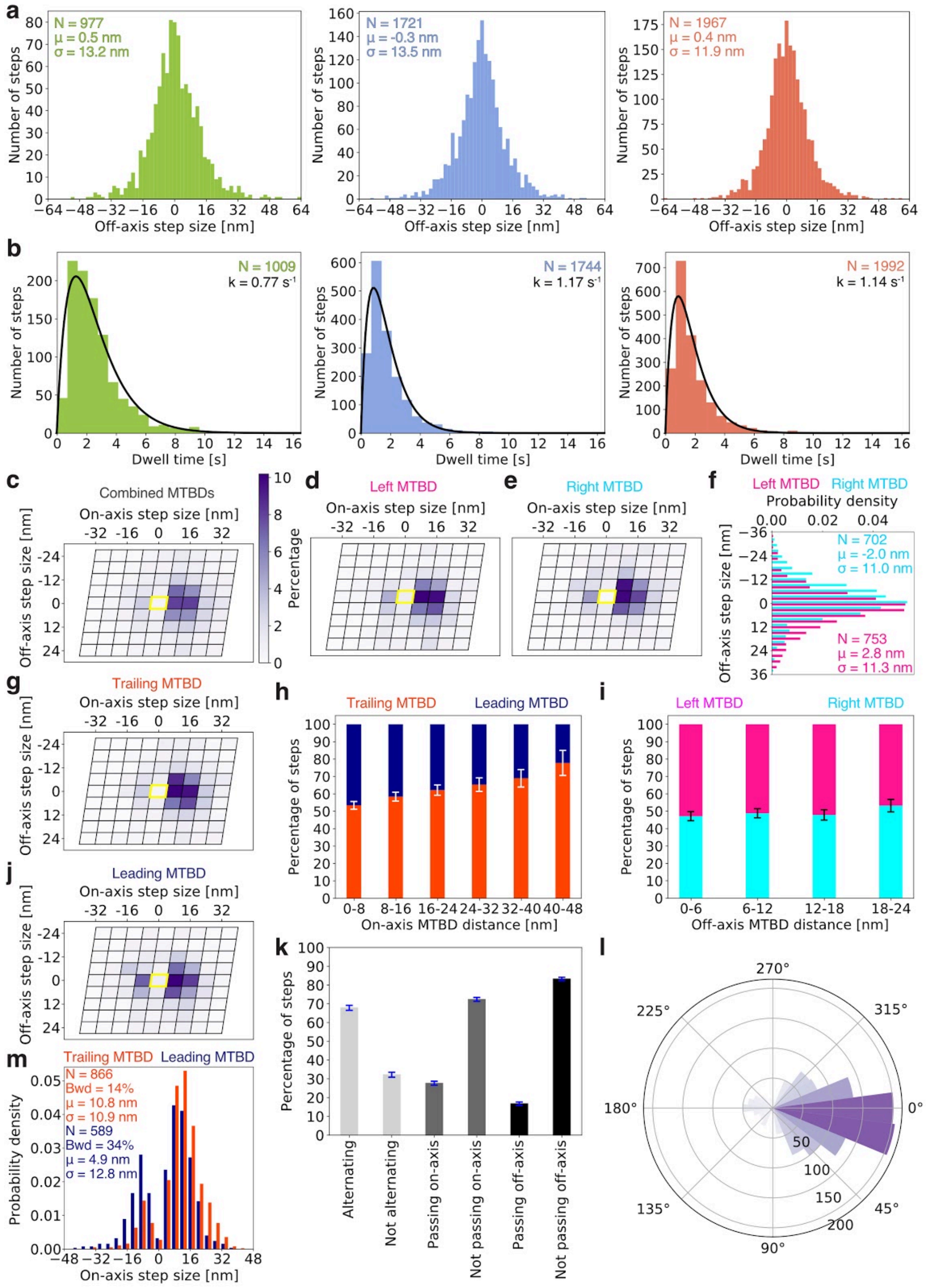


**Supplementary Figure 4 | Extended analysis of stepping parameters of both MTBDs and the AAA ring.** (**a**) Histogram of off-axis step sizes of dynein's AAA ring (green), the opposite MTBD (blue), and the associated MTBD (red). Number of steps detected (N), average step size (μ) and its standard deviation (σ) are given. (**b**) Histogram of the dwell times of dynein's AAA ring (green), the opposite MTBD (blue), and the associated MTBD (red). The black lines are fits of a convolution of two exponential functions with equal decay constants. (**c-e, g, j**) Heatmaps of the on- and off-axis step sizes of dynein's MTBDs mapped on the microtubule lattice. Here, each parallelogram represents a tubulin dimer consisting of one copy of $\alpha$ and $\beta$ tubulin. The yellow parallelogram represents the tubulin dimer at which the domain is located prior to the step. For each heatmap more than 1,000 steps are analyzed. (**c**) Step size distribution of all MTBDs regardless of relative position. (**d**) Step size distribution of all MTBDs that are to the left of the other MTBD prior to their step. (**e**) Step size distribution of all MTBDs that are to the right of the other MTBD prior to their step. (**f**) Histogram of off-axis step sizes of all MTBDs that are to the left (pink) or right (cyan) of the other MTBD prior to their step. (**g**) Step size distribution of all MTBDs that are trailing the other MTBD prior to their step. (**j**) Step size distribution of all MTBDs that are leading the other MTBD prior to their step. (**m**) Histogram of on-axis step sizes of all MTBDs that are trailing (orange) or leading (dark blue) the other MTBD prior to their step. (**f, m**) Number of steps detected (N), percentage of backward steps (Bwd), average step size (μ) and its standard deviation (σ) are given. (**h**) Probability of the trailing (orange) or leading (dark blue) MTBD to take the next step as a function of the inter-MTBD on-axis distance. (**i**) Probability of the left (pink) or right (cyan) MTBD to take the next step as a function of the inter-MTBD off-axis distance. (**k**) Percentage of steps for which the MTBDs alternate in stepping (light grey), for which the MTBDs passed each other when one of them took a step along the on-axis (medium grey) and along the off-axis (dark grey). (**l**) Angle histogram of the stepping angles. The stepping angle is defined as the angle between the 2D stepping vector and microtubule on-axis (See **Fig. 2 g**). A stepping angle between 0° and 90° refers to a forward-right step, a stepping angle between 90° and 180° refers to a backward-right step, a stepping angle between 180° and 270° refers to a backward-left step, and a stepping angle between 270° and 360° refers to a forward-left step. (**c-m**) The data for both MTBDs was combined. (**h, i, k**) The error bars show the bootstrapped standard error of the mean. (**h, i, k, l**) The total number of steps analyzed for these plots is N = 1,455.

The leading and trailing as well as the left and right MTBD prefer to step towards each other. For instance, a motor domain left of the other motor domain prefers to step towards the right (**d-f**) while a trailing motor is more likely to step forward than the leading motor (**c, g, j, m**). However, we only observe a bias for the trailing motor domain to take a next step with increasing inter-MTBD distance along the on-axis, while we do not see such bias for the left and right MTBD with increasing off-axis distance (**h, i**).

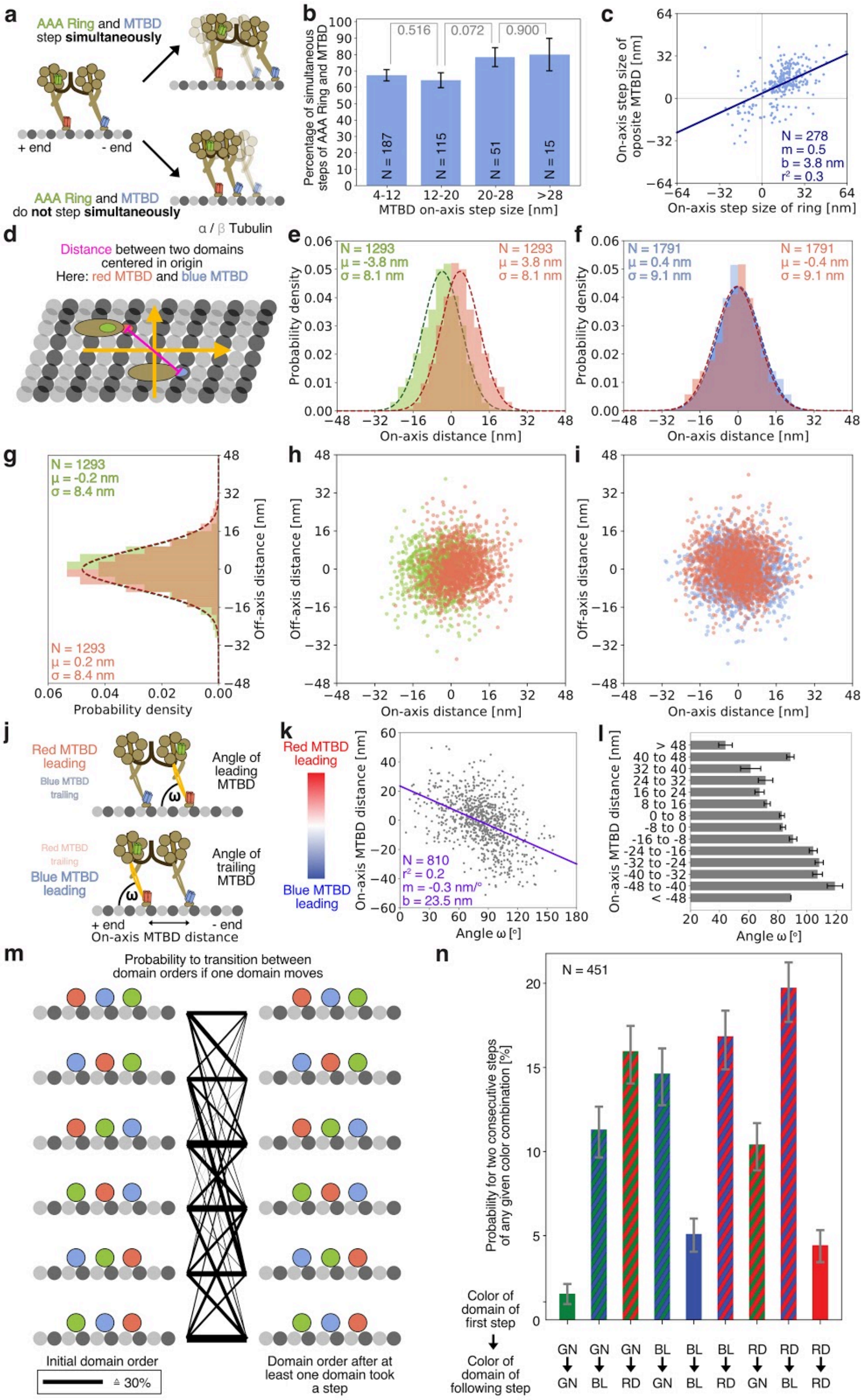

**Supplementary Figure 5 | Flexibility of stalk-microtubule angle.** (**a-c**) Analysis of simultaneous stepping of the AAA ring and MTBD on opposite motor domains. (**a**) The AAA ring and MTBD on opposite motor domains (green and blue, respectively) can either step simultaneously (top: both domains move along the on-axis) or not step simultaneously (bottom:

only MTBD moves while the AAA ring remains at the same on-axis position). (**b**) Histogram showing how often the AAA ring steps at the same time as the opposite MTBD (blue) as a function of the MTBD on-axis step size. N refers to the total number of steps for each condition. The error bars show the bootstrapped standard error of the mean. The p-values (grey) were calculated with a two-tailed z-Test. (**c**) Correlation of on-axis step sizes of the AAA ring and the opposite MTBD (blue) if they step at the same time. Each dot represents a single step. Blue line shows linear fit. N is the sample size. m is the slope. b is the y-intercept. $r^2$ is r squared value.

(**d-i**) Relative position of AAA ring and MTBD on the same motor domain and of both MTBDs on opposite motor domains. (**d**) Distance between two domains centered in origin. Here, the centroid position of either (**e, g, h**) AAA ring and MTBD or (**f, i**) both MTBDs are fixed in the origin and the distance of both domains relative to the centroid is measured. (**e, f**) Histogram of on-axis distances between (**e**) AAA ring (green) and the MTBD (red) of the same motor domain and (**f**) of on-axis distances between both MTBDs (blue and red). (**g**) Histogram of off-axis distances between AAA ring (green) and the MTBD (red) of the same motor domain. (**e, f, g**) Sample size (N), average distance (μ) and its standard deviation (σ) are given. (**h, i**) Scatter plot of the relative position between (**h**) AAA ring (green) and the MTBD (red) of the same motor domain and (**i**) both MTBDs (blue and red) on opposite motor domains. Here, the centroid position of either (**h**) AAA ring and MTBD or (**i**) both MTBDs is fixed in the origin and the position of both domains relative to the centroid is shown as a dot.

(**j-l**) Additional analysis for the stalk-microtubule angle as shown in **Figure 4**: The stalk-microtubule angle ω increases the more the motor domain is trailing (or less leading). (**j**) Schematic showing the definition of leading and trailing MTBD as well as the definition of the angle ω between stalk (orange) and on-axis microtubule lattice (small, grey circles). Note, the angle is only calculated for the motor domain for which the AAA ring (green) and the MTBD (red) are labeled. Top: Angle measurement for leading MTBD (red MTBD leading). Bottom: Angle measurement for trailing MTBD (blue MTBD leading). For more details see **Figure 4**. (**k**) Correlation between stalk-microtubule angles ω and inter-MTBD on-axis distances at the single-molecule level (grey dots). Here, a positive value refers to a state in which the red MTBD is leading and a negative value refers to a state in which the blue MTBD is leading. Purple line shows linear regression. N is the sample size. m is the slope. b is the y-intercept. $r^2$ is r squared value. (**l**) Same data as in **k** but binned into 8 nm bins (size of one tubulin dimer). Error bar shows standard error of the mean.

(**m, n**) Additional analysis for the transitions between the three-color domain orders as shown in **Figure 5**. (**m**) Probability to transition from one domain order to another for all six possible domain orders. A transition is counted if at least one domain took a step. The thickness of the black lines shows frequency. If no black line is drawn between two domain orders, this transition was not observed. (**n**) Frequency of two consecutive domain steps for any possible order. For instance, the left-most, green bar shows the frequency of a green-labeled AAA ring step followed by another green-labeled AAA ring step without movement of any other domain in between or at the same time along the on-axis. GN stands for green (AAA ring), BL stands for blue (opposite MTBD), and RD stands for red (associated MTBD). The error bars show the bootstrapped standard error of the mean.

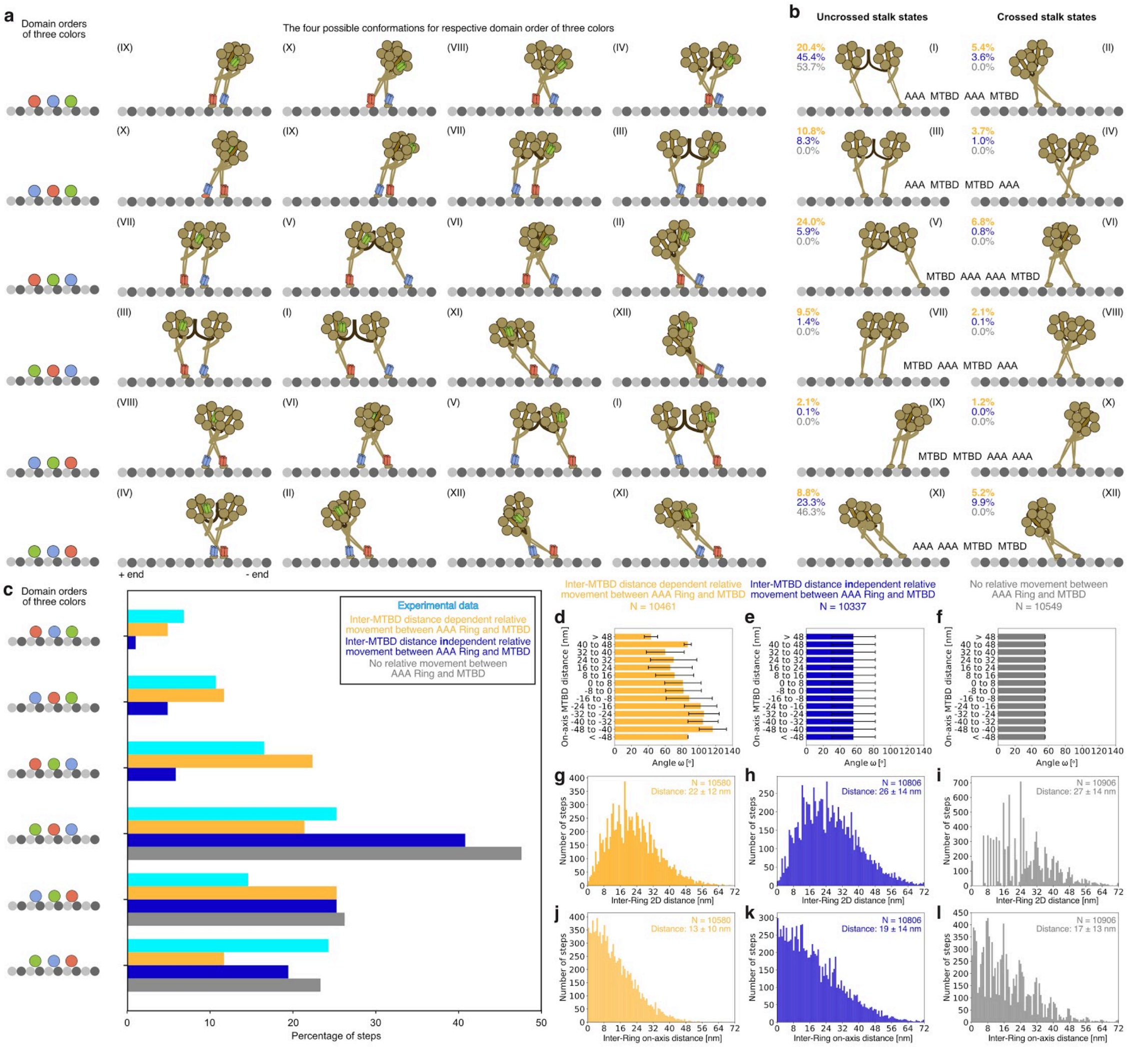


**Supplementary Figure 6 | Dynein adopts many conformations while moving along microtubules.** For this figure we only show representative conformations. Note that these are only based on the proximity of the respective domains (both AAA rings and both MTBDs) towards the microtubule minus end. Thus, any conformation shown here can have many other appearances for which the stalk-microtubule lattice angle (**Fig. 4**) or the inter-domain distances might be different. Moreover, we only show conformations for changes along the on-axis and are ignoring differences along the off-axis.

(**a**) Based on the three labels that we used in our three-color stepping experiments, we can find 6 possible domain orders (left-most section and **Fig. 5**). Each of these 6 domain orders can correspond to four of the 12 different dynein conformations shown in **b**, since the unlabeled AAA ring can be at any position. The Latin Numbers shown in the top left corner of each conformation correspond to the 12 conformations shown in **b.** Note that any of the 12 conformations in **b** is associated with two different domain orders of the three colors because the green AAA ring label with the associated red MTBD could also be on the other motor domain (dynein is a homodimer) yielding two possible domain orders of colors for any of the 12 dynein conformations. (**b**) Schematics showing all possible conformations dynein might be able to adopt if one allows both AAA rings to move relative to their respective MTBDs along the on-axis and ignores the absolute distance between these four domains but instead only focuses on the relative position towards the microtubule minus end. Since dynein is a homodimer, we can not distinguish the two motor domains and we end up with 12 possible conformations. These 12 conformations can be divided into 6 subcategories listed in the center. Within each subcategory, the two stalks can either cross (right) or not be crossed (left). For instance, there are two possible conformations for an order in which a AAA ring is furthest away from the microtubule minus end, lead by a MTBD, lead by another AAA ring, and finally lead by another MTBD (AAA-MTBD-AAA-MTBD). Using Monte Carlo simulations to simulate dynein stepping along microtubules as described in **Figure 6** and **Materials and Methods**, we quantified the occurrence of the 12 dynein conformations. We compared the occurrence of conformations for different regimes of stalk-microtubule on-axis angles (as introduced in **d-l**): for inter-MTBD distance dependent movement between AAA ring and MTBD as observed in this study (orange), for inter-MTBD distance independent movement between AAA ring and MTBD based on average values from cryo-electron microscopy studies[7] (blue), and for no relative movement between AAA ring and MTBD (grey). The percentages for all three cases and 12 possible conformations are listed in the top left.

The biggest difference for the 12 dynein conformations in **b** can be observed between the case where no relative movement between AAA ring and MTBD is allowed and the other two cases as the no movement case only allows for two of the twelve conformations. Moreover, the largest difference between the distance dependent relative movement (orange) and the distance independent relative movement (blue) is the occupancy among conformations 1, 5, 7, and 11 (counted from top left to bottom right).

(**c**) Comparison of frequency of three-color domain orders as introduced in **Figure 5**. Here, we are comparing the frequency of experimental observed three-color domain orders (cyan), to the frequency of three-color domain orders observed for Monte Carlo simulations for inter-MTBD distance dependent movement between AAA ring and MTBD as observed in this study (orange), for inter-MTBD distance independent movement between AAA ring and MTBD based on average values from cryo-electron microscopy studies[7] (blue), and for no relative movement between AAA ring and MTBD (grey). Note, that the experimental data is the same as in **Figure 5 b**.

Comparing the results of the frequencies of all possible six three-color domain orders from experimental and Monte Carlo simulated data (in **c**), we found that the Monte Carlo simulation, which used the inter-MTBD distance dependent movement between AAA ring and MTBD as observed in this study had the best agreement with the experimental data. We found the biggest difference between the experimentally observed three-color domain orders and the three-color domain orders observed for Monte Carlo simulations using an inter-MTBD distance dependent movement between AAA ring and MTBD when the associated, red-labeled MTBD was leading. However, overall the Monte Carlo simulated data and the experimental data agreed well, supporting the notion that the simulation recapitulates the experimental data well.

(**d-l**) Influence of relative movement between AAA ring and MTBD on inter-AAA ring distance. We used the Monte Carlo simulations to test how the relative movement between AAA ring and MTBD (see **Fig. 4**) influences the inter-AAA ring distance. Therefore we used different angles for the stalk-microtubule angle as input and looked at the inter-AAA ring distance as output. We compared (**d**) inter-MTBD distance dependent movement between AAA ring and MTBD as observed in this study, (**e**) inter-MTBD distance independent movement between AAA ring and MTBD based on average values from cryo-electron microscopy studies[7], and (**f**) no relative movement between AAA ring and MTBD. (**d-f**) Input values for the stalk-microtubule on-axis angle. The error bars show the standard deviation. (**g-i**) 2D inter-AAA ring distances for all three conditions. (**j-l**) On-axis inter-AAA ring distances for all three conditions. (**g-l**) Average and standard deviation of the distance values are provided. Note that the distance distribution in **g** and **i** for the flexible angle (yellow) and the fixed angle (grey) are the same data as in **Figure 6 c**.

We note that a previous study in which both AAA rings were labeled found a 2D inter-AAA ring distance of 18 ± 11 nm[8]. Thus, while the inter-MTBD distance dependent movement between AAA ring and MTBD (**d**) agrees well with the experimentally observed 2D inter-AAA ring distance, the other two cases (**e, f**) yield larger inter-AAA ring distances, showing that flexible motion between the AAA ring and MTBD might be necessary to maintain a closer proximity between both AAA rings.

(**b-l**) For all three cases we ran 100 Monte Carlo simulations (as described in **Figure 6** and **Materials and Methods**) for which we used a microtubule lattice with 13 protofilaments and a length of 79 tubulin dimers (~630 nm).

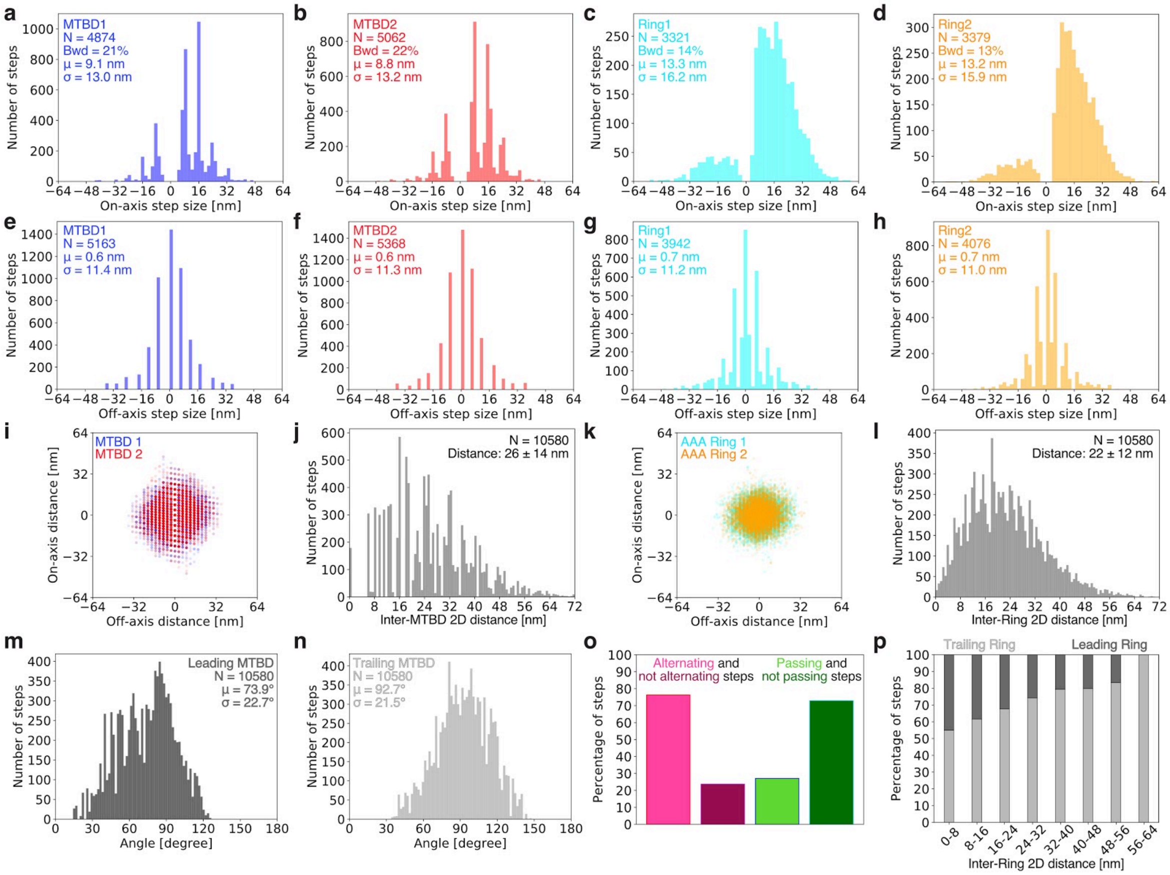


**Supplementary Figure 7 | Stepping analysis of AAA rings and MTBDs from Monte Carlo simulated data.** (**a-d**) Histogram of on-axis step sizes for (**a**) MTBD 1, (**b**) MTBD 2, (**c**) AAA ring 2, and (**d**) AAA ring 2. Here, the number of steps detected (N), percentage of backward steps (Bwd), average step size (μ) and its standard deviation (σ) are given. (**e-h**) Histogram of off-axis step sizes for (**e**) MTBD 1, (**f**) MTBD 2, (**g**) AAA ring 2, and (**h**) AAA ring 2. Here, the number of steps detected (N), average step size (μ) and its standard deviation (σ) are given. (**a-h**) Note that the step sizes for the MTBDs are discrete and that the step sizes for the AAA rings are more continuous since only the MTBDs follow the well defined microtubule lattice. (**i, k**) Scatter plot of the relative position between (**i**) both MTBDs and (**k**) both AAA rings. Here, the centroid position of either (**i**) MTBDs or (**k**) AAA rings is fixed in the origin and the position of the domains relative to the centroid is shown as a dot. (**j**) Histogram of the inter-MTBD 2D distance. (**l**) Histogram of the inter-AAA ring 2D distance. (**i-l**) Overall, the AAA ring spans a smaller area than the MTBDs and the average distance between both MTBDs is larger than the average distance between the AAA rings. This agrees well with the fact that the two AAA rings are held together by the tail and that the MTBDs likely can move relative to their respective

AAA ring and therefore explore a larger area. (**m**) Histogram of angles between dynein's stalk and the microtubule lattice for the motor domain for which the MTBD is leading. (**n**) Histogram of angles between dynein's stalk and the microtubule lattice for the motor domain for which the MTBD is trailing. (**o**) Percentage of steps for which the AAA rings alternate in stepping (pink) or do not alternate in stepping (purple) and for which the AAA rings passed each other when one of them took a step along the on-axis (light green) or did not pass each other (dark green). (**p**) Probability of the trailing (light grey) or leading (dark grey) AAA ring to take the next step as a function of the inter-AAA ring on-axis distance. (**o, p**) These data agree well with previous observations for which both AAA rings were labeled while dynein was stepping along microtubules[8] showing that our Monte Carlo simulation recapitulates previous observations. (**a-p**) All this data is from 100 Monte Carlo simulations (as described in **Figure 6** and **Materials and Methods**) for which we used a microtubule lattice with 13 protofilaments and a length of 79 tubulin dimers (~630 nm).

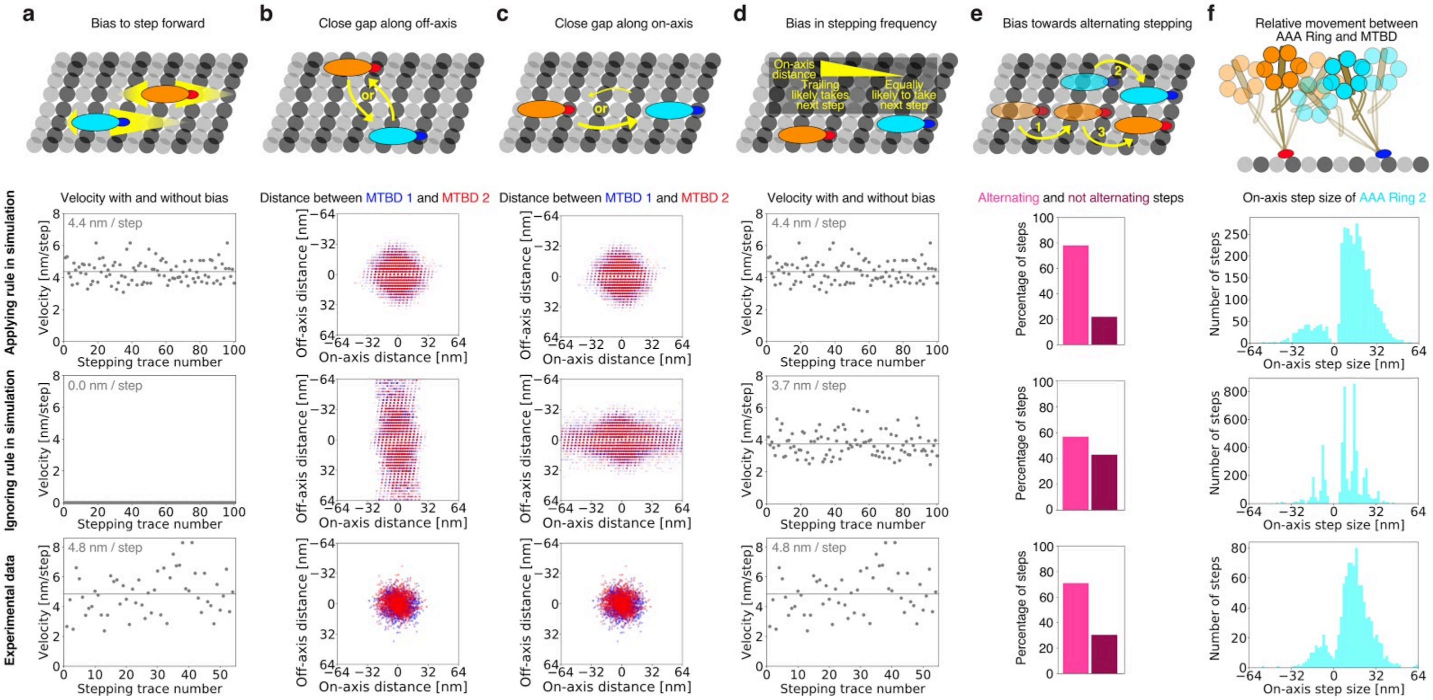


**Supplementary Figure 8 | Effects of Monte Carlo simulation rules on dynein motility.** (**a-f**) Schematics on top represent requirements for dynein motility. The graphs below show an example of what happens to dynein motility if the requirement is applied (top graph) or ignored (middle graph) during the simulation as well as the corresponding experimental data (bottom graph). (**a**) If the bias to step forward is ignored, dyneins performs random walks with no positional net gain. (**b**) If the bias of the stepping direction along the off-axis is ignored, both motor domains often drift apart along the off-axis. (**c**) If the bias of the stepping direction along the on-axis is ignored, both motor domains often drift apart along the on-axis. (**d**) If the bias in stepping frequency of the leading and trailing motor domain is ignored, dynein will move more slowly towards the microtubule minus end. (**e**) If no bias towards alternating stepping is encoded, dynein will have a lower tendency to alternate. (**f**) If there is no relative movement between AAA ring and MTBD, the AAA ring will step exactly as the MTBD and follow the microtubule lattice. Moreover, the inter-AAA ring distance will change (**Supplementary Fig. 6 d-l**). Note, that the experimental data in **b** and **c** is the same as in **Supplementary Figure 5 i**. Note, that the experimental data in **e** is the same as in **Supplementary Figure 4 k**. Note, that the experimental data in **f** is the same as in **Figure 2 a and b**.

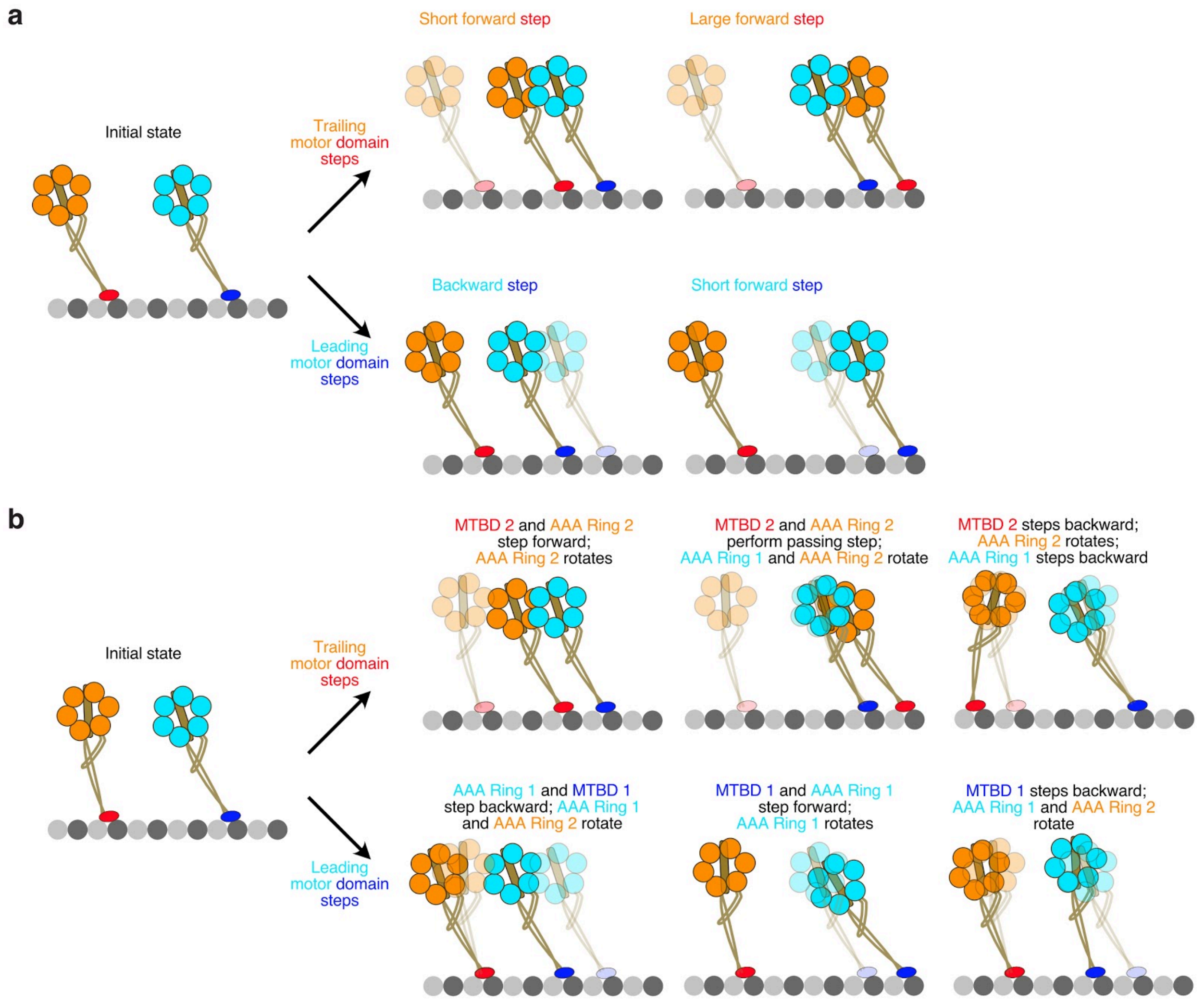


**Supplementary Figure 9 | Models for dynein stepping.** (**a**) Hypothetical model for dynein stepping not accounting for flexible movement between AAA rings and MTBDs. Left: Initial state in which both motor domains have the same angle between stalk and microtubule on-axis. Top: Potential steps that can be taken by the trailing motor domain - a short forward step during which the stepping motor domain does not pass the other motor domain or a larger forward step during which the stepping motor domain passes the other motor domain. Bottom: Potential steps that can be taken by the leading motor domain - a short backward step or a short forward step. (**b**) Another model for dynein stepping that does take into account the observed flexibility between AAA ring and MTBD of the same motor domain. Left: Initial state in which both motor domains have a slightly different angle between stalk and microtubule (see **Fig. 4 f**). Top: Potential steps that can be taken by the trailing motor domain. Bottom: Potential steps that can be taken by the leading motor domain. (**a, b**) Opaque motor domains show initial state. When we compared the stepping motions in which the relative movement between AAA ring and MTBD is not taken into account (**a**) or in which it is allowed (**b**), we found that the rigid model can not explain the experimentally observed conformational variety (**Fig. 5, 7**). In particular, the stepping motions for the trailing as well as the leading motor domain shown on the farright of **b** are different than previously described. Here, the MTBD can for instance take a small backward step while the AAA ring does not seem to translocate along the microtubule on-axis and only rotates.

### Supplementary Information Tables

| Name | Sequence and modification | Vendor |
|---|---|---|
| FC_SC_01_ATTO488 | /5ATTO488N/ATGAGGTGTATGTGTAGAGTGATGGATGTAGT/3ATTO488N/ | IDT |
| FC_SC_02_ATTO488 | /5ATTO488N/AGGATGAGTGAGAGTGAGATGAGAGTAGATGT/3ATTO488N/ | IDT |
| FC_St_02_ATTO488 | /5ATTO488N/CACTCTCACACCTCATACATCTACCATCACTC/3ATTO488N/ | IDT |
| FC_SC_01_Cy3N | /5Cy3N/ATGAGGTGTATGTGTAGAGTGATGGATGTAGT/3Cy3N/ | IDT |
| FC_SC_02_Cy3N | /5Cy3N/AGGATGAGTGAGAGTGAGATGAGAGTAGATGT/3Cy3N/ | IDT |
| FC_St_02_Cy3N | /5Cy3N/CACTCTCACACCTCATACATCTACCATCACTC/3Cy3N/ | IDT |
| FC_SC_01_ATTO647N | /5ATTO647NN/ATGAGGTGTATGTGTAGAGTGATGGATGTAGT/3ATTO647NN/ | IDT |
| FC_SC_02_ATTO647N | /5ATTO647NN/AGGATGAGTGAGAGTGAGATGAGAGTAGATGT/3ATTO647NN/ | IDT |
| FC_St_02_ATTO647N | /5ATTO647NN/CACTCTCACACCTCATACATCTACCATCACTC/3ATTO647NN/ | IDT |
| FC_St_01_HALO | Halotag Ligand (O2) TACACATACTCATCCTACTACATCTCTCATCT | Biomers |
| FC_St_01_CoA | Coenzyme A (CoA) TACACATACTCATCCTACTACATCTCTCATCT | Biomers |

**Supplementary Table 1 | Sequences for assembly of DNA FluoroCubes.** The sequences used for the DNA FluoroCubes are the same as previously published[4]. For each FluoroCube four oligonucleotides are required: SC_01, SC_02, St_01, and St_02. Here, we used three fluorescently labeled oligonucleotides (SC_01, SC_02, and St_02) and one functionally tagged oligonucleotide (St_01, grey part of table) to assemble FluoroCubes.

| Name | Sequence | Vendor |
|---|---|---|
| Core_001 | GTAAATTGTGTAGAGCACATTTG | IDT |
| Core_002 | CGGCCGCCAGTTCGGGCCTCGGA | IDT |
| Core_003 | GCTGACCATCAATAGCATGACAA | IDT |
| Core_004 | AAATAATATAATAACCTGTAAAT | IDT |
| Core_005 | ATAGTACATAACCTGGATAAGAG | IDT |
| Core_006 | TGCTCTGATAAAGACCTTCAACA | IDT |
| Core_007 | GTTGAAGGAATTACAACAACACT | IDT |
| Core_008 | AAACACTGCCTAGAATAGGTTTC | IDT |
| Core_009 | AGAATGATAGCACACCACCTGTA | IDT |
| Core_010 | CTATCGAGCAAGACTCCTTTAGA | IDT |
| Core_011 | GTAGGAACAAGCTTAAATCAAGT | IDT |
| Core_012 | ATACCCTTCTGAAACGCTCAATC | IDT |
| Core_013 | TCGCCTTTACATATTTAACCTGA | IDT |
| Core_014 | ATCTGACCTCAATTTAGAATAAA | IDT |
| Core_015 | AGAATCCGAGTAAAAGATTAACA | IDT |
| Core_016 | CTTTCATAAATCGTGAGCTAACTCAC | IDT |
| Core_017 | GGGTCTTTCTGCTCTCGCACTCAATC | IDT |
| Core_018 | GTGAGCTTGTAGATCGAAACGTACAG | IDT |
| Core_019 | TAAATGGGGGTGAGTCGCGTCTGGCC | IDT |
| Core_020 | TCAACTTTTTGCGGTATGACCCTGTA | IDT |
| Core_021 | AAGCGGATTAGGAACGAAAGACTTCA | IDT |
| Core_022 | CGTTGACCAAAGTAATGCGATTTTAA | IDT |
| Core_023 | GCACCAAAGCGGAGTCCATTAAACGG | IDT |
| Core_024 | ATCTAGAAACAGTGGTAGCATTCCAC | IDT |
| Core_025 | GTTCCGGAAGGCCGTTAAAGCCAGAA | IDT |
| Core_026 | AGGTGGACCACAAGGGGAGGGAAGGT | IDT |
| Core_027 | GAAGCAAGGTATTAAATAGCAGCCTT | IDT |
| Core_028 | AAAAAAACTTTTTCCGCGCCTGTTTA | IDT |
| Core_029 | AACCTAAACGTCAGAATTTATCAAAA | IDT |
| Core_030 | ATGGACAAATGAAACAATATAATCCT | IDT |
| Core_031 | TCTGTTGAGCCTTACAGACGATCCAGCG | IDT |
| Core_032 | GTTAACGGAACGGGAAAAGCCGCACAGG | IDT |
| Core_033 | TTCTGTACGCCACTTCAGGAAGATCGCA | IDT |
| Core_034 | CTACATGTACCCCATAAATTAATGCCGG | IDT |
| Core_035 | TAACATTTTCATGATGCAACTAAAGTAC | IDT |
| Core_036 | GATAAACTGCGGATAAGAGCAACACTAT | IDT |
| Core_037 | ATAAGAATCTTGCCCATGTTACTTAGCC | IDT |

| Name | Sequence | Vendor |
|---|---|---|
| Core_038 | TGTATACGCATAAAAAATCTCCAAAAAA | IDT |
| Core_039 | TTAGGGTACTCACCTGAAAGTATTAAGA | IDT |
| Core_040 | TCGGCTCCCTCATCCAAGTTTGCCTTTA | IDT |
| Core_041 | GAACATATGTTAAATTACCGAAGCCCTT | IDT |
| Core_042 | GAAGGTGCTATTAGAATCATTACCGCGC | IDT |
| Core_043 | CCGGAGCTTAATATGACCGTGTGATAAA | IDT |
| Core_044 | GCGAATGAATTACTTGATTGCTTTGAAT | IDT |
| Core_045 | GGTTATTTACAAGAGTCAGTTGGCAAAT | IDT |
| Core_046 | TCCCAGTCGGGTACATGCGCCTGTGCAC | IDT |
| Core_047 | GTGCCCACAGCGTGACATGAAGGGTAAA | IDT |
| Core_048 | GCATTCCGCTGCAATTTTCCGGCACCGC | IDT |
| Core_049 | TTAGCCCCCCCAAACTTTTTTGAGAGAT | IDT |
| Core_050 | TTACCTGGGTGGCAGGTTTCATTCCATA | IDT |
| Core_051 | ACGAGATTGAATCCGATTACCAGACGAC | IDT |
| Core_052 | GAACGTTTACCCAAATAGACGGTCAATC | IDT |
| Core_053 | CCCAATTGAGGCTTACGGAGCCTTTAAT | IDT |
| Core_054 | GACAGAACCTCAGAAATCAAGAGAAGGA | IDT |
| Core_055 | AATAGCACGCCACCGAAGCGCGTTTTCA | IDT |
| Core_056 | AGCCAATACATAAATTAAGCAGATAGCC | IDT |
| Core_057 | AAAAGACTATCCTGTTAAATCAGATATA | IDT |
| Core_058 | CTTAGCACAACGCCAATAAGAATAAACA | IDT |
| Core_059 | AAACCAATACATAAACAATCGCGCAGAG | IDT |
| Core_060 | GTTTTTTATGGGATAGACGTTTAGCAAG | IDT |
| Core_061 | GTAAGATGTAAATCCATGGAATTGAGGAA | IDT |
| Core_062 | GGCCGCGGGTGCTGCGGCTTACACTGCGCCAG | IDT |
| Core_063 | ACGCGAAACGATGCTGATTAGCGGATGCTGATT | IDT |
| Core_064 | TTCGCATGTGCCGGAAACCATGGGATAGCGGTC | IDT |
| Core_065 | GTCTGTAAAGGCTATCAGGTGTAGGTTGATGGT | IDT |
| Core_066 | TAGATTAGTTGATTCCCAACTTTTGAAGAATAG | IDT |
| Core_067 | CAAAAATAAACCAAAATAGAAGATTCCAAGAGT | IDT |
| Core_068 | GAGGAGTGGAACCGAACTGGATTATAGCGAAAA | IDT |
| Core_069 | GGCGGCCATTAGCGGGGTTCGGGGTCACTAAAT | IDT |
| Core_070 | GCGTTCCATTTTCGGTCATTAGCACCACGGGGA | IDT |
| Core_071 | ACGCAAGAAGTTACCAGAACTAATATAGCGAAA | IDT |
| Core_072 | GGCGTGTCTTATCCGGTATCCTTATCCACGCTG | IDT |
| Core_073 | TTAGTGGATCATAATTACTGACAAAGGCTACAG | IDT |
| Core_074 | AAGAATTTTATTCATTTCAGCGTAGAACGTGCT | IDT |
| Core_075 | CAACTACTCTAAAATATCTACGCTGAGGCCGAT | IDT |

| Name | Sequence | Vendor |
|---|---|---|
| Core_076 | CGGTTGGGCTGCCGGACTTGTGCAACCGCAAGAAT | IDT |
| Core_077 | AACGTGCATGCTGGCGAAAGGCGCCAGGGTTTTCC | IDT |
| Core_078 | TGGTCAAATCGGTTGATAATCGCAAATATTTAAAT | IDT |
| Core_079 | AAAATTGCTTTGGGGCGCGAGGTAGCATTAACATC | IDT |
| Core_080 | AAAATGCAGAATCGTCATAAAAGTTCAGAAAACGA | IDT |
| Core_081 | GAATCATCGACAAGAACCGGACTCATTCAGTGAAT | IDT |
| Core_082 | AGAACAACTACCGATATATTCCTTTTGCGGGATCG | IDT |
| Core_083 | GGTAATGCCGGAGGTTTAGTAGCCACCCTCAGAGC | IDT |
| Core_084 | CCGAAACCAGAGCCGCCACCCAGAGCCGCCACCAG | IDT |
| Core_085 | TTCAATAATGCAAACGTAGAAAAACGCAAAGACAC | IDT |
| Core_086 | ACCTCATCGTTGCACCCAGCTGAGCGTCTTTCCAG | IDT |
| Core_087 | AATAATTTCTGAGAATCGCCAAGGCATTTTCGAGC | IDT |
| Core_088 | ATACAGTACCCTTTTTTAATGATAACCTTGCTTCT | IDT |
| Core_089 | GCCCTTGCTACAATTCGACAAATTTTAAAAGTTTG | IDT |
| Core_090 | GAACCGTTGTTACATTGGCAGACATTCTGGCCAAC | IDT |
| Core_091 | GGTGGTTTTTCTTTGTCATAAGTAATGGGGTGCCA | IDT |
| Core_092 | GCCCTTCACCGCCTAGAGACGGTGAAGGACGGCCA | IDT |
| Core_093 | CACGCTGGTTTGCCGCGGATTGTCGGATAAAATTC | IDT |
| Core_094 | GGTTCCGAAATCGGATGCCTGAGAACCCCAAAGAA | IDT |
| Core_095 | CCCGAGATAGGGTTACCTTTACTCCAACCAGGTCT | IDT |
| Core_096 | CCACTATTAAAGAATTATTACAATAAAACACCAGA | IDT |
| Core_097 | ACCGTCTATCAGGGCTTTGACCAAAAGAAGCATCG | IDT |
| Core_098 | CGGAACCCTAAAGGGTAATAACTTTTGACAGCATT | IDT |
| Core_099 | AAGCCGGCGAACGTAGCAAAACTTGAGCCACAATC | IDT |
| Core_100 | GGAGCGGGCGCTAGAGGGTAAAACTGAAAATAAAC | IDT |
| Core_101 | CGCGTAACCACCACAATCGGCTACGAGCTACCGAC | IDT |
| Core_102 | GGCGCGTACTATGGAAATCCATATAACTATTTTCC | IDT |
| Core_103 | TTCCTCGTTAGAATGAAATAAATCAAAAAACAAAG | IDT |
| Core_104 | TAAAGGGATTTTAGCCTGCAAGGTGAGGTGAAAGC | IDT |
| Core_105 | GTAATGGATCCCGCCTAATGAGTTAAGTGTAAAGCCTG | IDT |
| Core_106 | GAATTCCCGGTTTATCAGCAACAACTATCACCCAAATCAA | IDT |
| Core_107 | TCACTGCAATACCTCAATCGTCTGAACAACAGGAAAAACG | IDT |
| Core_108 | CAGCAAGAACGTTTGCAGGCGCTTATCCAGCATCAGCGGG | IDT |
| Core_109 | GGTAAGGGATGTGGAAACAATCGGGGGGAACGGATAACCT | IDT |
| Core_110 | TATAAAGAAAAGATCAAAAATAATAATTAACCAATAGGAA | IDT |
| Core_111 | TAGTACTGAAAATACCAAAAACATATATAAAGCTAAATCG | IDT |
| Core_112 | TAAACTATTCATTAAGAGGAAGCCTAGGATTGCATCAAAA | IDT |
| Core_113 | AGCTGTATTCATGTGAATTACCTTCAATTTCAACTTTAAT | IDT |

| Name | Sequence | Vendor |
|---|---|---|
| Core_114 | GGCCGGGTCGCTCATGAGGAAGTTTGGAGGACTAAAGACT | IDT |
| Core_115 | GAACCCCGCCACACTACAACGCCTCCGTTTCGTCACCAGT | IDT |
| Core_116 | CCCTCTCAGAACAATAAATCCTCAGACCTTGATATTCACA | IDT |
| Core_117 | AAAAGAATACATTCAACCGATTGAAAAAGACAAAAGGGCG | IDT |
| Core_118 | CTAACACAATTTGTCAAAAATGAAAAAACGATTTTTTGTT | IDT |
| Core_119 | GGCAGTATTTAAGCTAATGCAGAAAACAATAAACAACATG | IDT |
| Core_120 | AGTGAGAAACAGGAGTCAATAGTGATAGATTAAGACGCTG | IDT |
| Core_121 | TATTACTCGTATATGGCAATTCATAAATTCCTGATTATCA | IDT |
| Core_122 | GGTGTTTGCCGCCAGCAGTGTAAAGGACTGTTGCCCTGCG | IDT |
| Core_123 | CCACGGCGCATCGTAACCGGATAGCTAGATAGACTTTCTC | IDT |
| Core_124 | ATTTTAGGCCGGAGACATTTCTCCGTGTGAGCGAGTAACA | IDT |
| Core_125 | AGAGCGGCTTAGAGCTTGGTCATATAGCAAGGATAAAAAT | IDT |
| Core_126 | AAAGCCCACATTCAACTATAGGTCAGAACCAGACCGGAAG | IDT |
| Core_127 | GTTTAACGGAGATTTGTAACGAACTAGGAAGAAAAATCTA | IDT |
| Core_128 | GCTTTAGAATAGAAAGGACATACACTAACCTAAAACGAAA | IDT |
| Core_129 | ACTGACGTATAAACAGTGAAGTAAATAAGTTTTGTCGTCT | IDT |
| Core_130 | ATTGGAACGTCACCAATGCTGATACAAGTAAGCGTCATAC | IDT |
| Core_131 | CGCCATTGAGTTAAGCCGTCATTTGGAATTATCACCGTCA | IDT |
| Core_132 | TAAGACCAAGTACCGCAAACACCCTGGCATTAGACGGGAG | IDT |
| Core_133 | GACGAATATATTTTAGTAGATGTAGATAATATCCCATCCT | IDT |
| Core_134 | AGCTTGAATATACAGTATAATATGTACCGGCTTAGGTTGG | IDT |
| Core_135 | ATCATTCTAAAGCATCAGGTTATTTGAGGGTTAGAACCTA | IDT |
| Core_136 | TTGAATCACGCAAATTACCCGGTCAGAGCAGAAGATAAAA | IDT |
| Core_137 | CAACACAAACTTAAATTTCTCCTCATGTCCGTTTTTTCGT | IDT |
| Core_138 | CAGCAAGGTCACGTTGGCACTTCGCTCATTCAGGCTGCGC | IDT |
| Core_139 | CTGTTAAAGATTCAAAAGATCAATCAGAGCAAACAAGAGA | IDT |
| Core_140 | TCAAATAAGAGGTCATTTTTTAGCTATTAGTTTGACCATT | IDT |
| Core_141 | TGGAAATCAGTTGAGATTTGCGTCCAGAAGTTTTGCCAGA | IDT |
| Core_142 | AAAGGCCAAGCGCGAAAGGATCAAGACAGATGAACGGTGT | IDT |
| Core_143 | GAACCTTCAACAGTTTCAACCATCGCTCTTAAACAGCTTG | IDT |
| Core_144 | AAAGCAGTGCCTTGAGTATTGTATCAATAAGTGCCGTCGA | IDT |
| Core_145 | GCTTGATTACCATTAGCAGGGAACCGTGCCATCTTTTCAT | IDT |
| Core_146 | AAGAACAGAGAGATAACATATTACGCATAATAACGGAATA | IDT |
| Core_147 | GCGGTATTCCAAGAACGCAAAGATTATTTAGCGAACCTCC | IDT |
| Core_148 | GCGCCAACGCGAGAAAACCAACAGTAATCATATGCGTTAT | IDT |
| Core_149 | GTATATTTTCAGGTTTATGAATTTCAGATGATGAAACAAA | IDT |
| Core_150 | CAGGAGAGCCAGCAGCACGGTATTAGAATAGATTAGAGCC | IDT |
| Core_151 | ATTAATGACATCCCCAAGCCTCCGGCCACATCGACCGACCGT | IDT |

| Name | Sequence | Vendor |
| --- | --- | --- |
| Core_152 | CGCCGTGCAGAAACGCGGGCGATCGGTGTGAGGGGGTTGTAA | IDT |
| Core_153 | CGCCACCGACCGTAAGTAATCGTAAAACTATGATAGGTATTT | IDT |
| Core_154 | TTCCTTTAGTAATGTCCAAATGGTCAATGCTGTAGAACAGGC | IDT |
| Core_155 | ATACTAAATTGCTCTTAAAATGTTTAGAGCCAAAATCATCAA | IDT |
| Core_156 | AATATTTAGGTAGACGCATAGGCTGGCTTTGTGTCCCGACGA | IDT |
| Core_157 | GAACTGGCCCCAGCACGCGCCGACAATGGCGAATAATGCGAA | IDT |
| Core_158 | GTAAACCTTTGCTATTAAGTATAGCCCGTTTCGGAGCTTTCA | IDT |
| Core_159 | AGACAGGGTTTTAATTCGGAACCAGAGCGCACCGTACAGCCG | IDT |
| Core_160 | TGGAAGATCACCAGAGGGCATGATTAAGAAACAATCTTTATT | IDT |
| Core_161 | AAATATTTTGAGCGGGGGTTTTGAAGCCAAGCCGTGGCAGTT | IDT |
| Core_162 | TACAGTTTGTCTTTTCCAGTATAAAGCCCCTAAATAAAATAT | IDT |
| Core_163 | TCAACTAATCGCAAAGAAAATTAATTACCGGGAGAAATATTA | IDT |
| Core_164 | TCATAATAGAAATTATTACATTTGAGGAATATCAAATATTTT | IDT |
| Core_165 | GATTGGACAGTGCCTTCTACATTTTGACGTTCTTTGTTCTTCT | IDT |
| Core_166 | ACCAGCACGCGTGCCCAGCGGTGA | IDT |
| Core_167 | ACCGGTGCCCCCTGCATCCTGCAGCTGTTCTATCGGCCAACGCA | IDT |
| Core_168 | GTCATAGCTGTTTCCTGTGTGAA | IDT |
| Core_169 | AAATTGTTGCCGGGTCCTCACAGTTGACATG | IDT |
| Core_170 | ACACACAACATACGAGCCGGA | IDT |
| Core_171 | AGCATAAACGGCATCAGATATTGCATTACAGTCGGGAAACCTGTA | IDT |
| Core_172 | ACGTGCCAGCCCGCTCACAATTCA | IDT |
| Core_173 | TCAGCGATCGCGTCTTTTCACGGTCATACCGGGGGTTTCTGA | IDT |
| Core_174 | AGCGGGGAGAGGCGGTTTGCGTATTGGGGTGTGT | IDT |
| Core_175 | ATACCGCCAGCCATTGATTCCAGAACAATATA | IDT |
| Core_176 | ACGGCCTTGCTGGTAATAGGATTATTATGCCTGA | IDT |
| Core_177 | GTAGAAGAACTCAAACTATA | IDT |
| Core_178 | AACCAGTAATAAAAGGGATTCACCAGTCACACGA | IDT |
| Core_179 | AAATGCGCGAACTGCATGGCTATTAGTCTTTA | IDT |
| Core_180 | AACCACCTAGTCTGTCATAGCCCTAAAACATCGA | IDT |
| Core_181 | ACCATTAAAAATACCGAACG | IDT |
| Core_182 | AATAATCAGTGAGGCCACCTGAGAAGTGTTTTTA | IDT |
| Handle site helix0, #0 | CTCGTCGCTGGCGAATGCGGCG | IDT |
| Handle site helix0, #1 | AACTGTTGGGAACGTTCCGGCAA | IDT |
| Handle site helix0, #2 | ATCGATGAACGGGCAAAGCGCCA | IDT |
| Handle site helix0, #3 | AGATACATTTCGATTGCCTGAGA | IDT |
| Handle site helix0, #4 | GGGGGTAATAGTCTGCGAACGAG | IDT |
| Handle site helix0, #5 | ACAGACCAGGCGAGAGGCTTTTG | IDT |
| Handle site helix0, #6 -ATTO488 | ATACCGATAGTTCAACTTTGAAATT/3ATTO488N/ | IDT |

| Name | Sequence | Vendor |
|---|---|---|
| Handle site helix0, #7 -Cy3 | GAGGGTTGATATGCTTTCGAGGTTT/3Cy3Sp/ | IDT |
| Handle site helix0, #8 -ATTO647N | AATCAAAATCACGCTCAGTACCATT/3ATTO647NN/ | IDT |
| Handle site helix0, #9 | CCCAAAAGAACTCCCCCTTATTA | IDT |
| Handle site helix0, #10 | CGACTTGCGGGAAAACCGAGGAA | IDT |
| Handle site helix0, #11 | ACAAATTCTTACTAAGAACGCGA | IDT |
| Handle site helix0, #12 | CATCAAGAAAACAAAAAGCCTGT | IDT |
| Handle site helix0, #13 | GTCAATAGATAATACCTGAGCAA | IDT |
| Handle site helix0, #14 | CTCATGGAAATACAGGAGCACTAA | IDT |
| Handle site helix3, #0 | CAGTGTCACTGCAAAAAAATCCC | IDT |
| Handle site helix3, #1 | CGGCCTTTAGTGGACGACAGTAT | IDT |
| Handle site helix3, #2 | CTCCAGCCAGCTCAACCGTTCTA | IDT |
| Handle site helix3, #3 | AGAGGGTAGCTACAACATGTTTT | IDT |
| Handle site helix3, #4 | GGTGTCTGGAAGAATTACGAGGC | IDT |
| Handle site helix3, #5 | CATAACCCTCGTAATCCGCGACC | IDT |
| Handle site helix3, #6 | GGAACGAGGCGCAATTTTTTCAC | IDT |
| Handle site helix3, #7 | AAGGCTCCAAAACTATTATTCTG | IDT |
| Handle site helix3, #8 | GGCTGAGACTCCTCAGTAGCGAC | IDT |
| Handle site helix3, #9 | GCGTCAGACTGTAATAGCAATAG | IDT |
| Handle site helix3, #10 | TTTAAGAAAAGTTTATTTTCATC | IDT |
| Handle site helix3, #11 | CCAATAGCAAGCAATGGTTTGAA | IDT |
| Handle site helix3, #12 | TAAGGCGTTAAACAATAACGGAT | IDT |
| Handle site helix3, #13 | ACCAAGTTACAACCTCAATCAAT | IDT |
| Handle site helix3, #14 | CAACAGTTGAAATAGTAATAACA | IDT |
| Handle site helix4, #0 | GCCAACGGCAGCAGCTCGAATTC | IDT |
| Handle site helix4, #1 | CAGTCACGACGTGCTGGTCTGGT | IDT |
| Handle site helix4, #2 | TGTAAACGTTAACGATTAAGTTG | IDT |
| Handle site helix4, #3 | CAATAAATCATACAGGAAGATTG | IDT |
| Handle site helix4, #4 | GAATGACCATAAAATTCTACTAA | IDT |
| Handle site helix4, #5 | AAGGCTTGCCCTCTCAAATGCTT | IDT |
| Handle site helix4, #6 | TCACCCTCAGCACAACGTAACAA | IDT |
| Handle site helix4, #7 | CACCACCCTCATAGGGAGTTAAA | IDT |
| Handle site helix4, #8 | AACCACCACCAGCGCCACCCTCA | IDT |
| Handle site helix4, #9 | CACGGAATAAGTCAGAGCCACCA | IDT |
| Handle site helix4, #10 | AGCCTAATTTGCTGGCAACATAT | IDT |
| Handle site helix4, #11 | CAGTAATAAGAGTCTTACCAACG | IDT |
| Handle site helix4, #12 | GTAAATCGTCGCCATGTAATTTA | IDT |
| Handle site helix4, #13 | AGTAACATTATCCAATATATGTG | IDT |
| Handle site helix4, #14 | AGAGATAGAACCTGCCCGAACGT | IDT |

| Name | Sequence | Vendor |
|---|---|---|
| Handle site helix7, #0 -Bio | GGCGCAACCAGCTTACGGCTGGA TTT/3Bio/ | IDT |
| Handle site helix7, #1 | GTAAGCTTTCAGAGGTGGAGCCG | IDT |
| Handle site helix7, #2 | CAAAATTTTTGTTAAATCAGCTC | IDT |
| Handle site helix7, #3 | CGAAAATTAAGCAATAAAGCCTC | IDT |
| Handle site helix7, #4 | GTCTGACTATTATAGTCAGAAGC | IDT |
| Handle site helix7, #5 -Bio | AGTAGTAAATTGGGCTTGAGATG TTT/3Bio/ | IDT |
| Handle site helix7, #6 -Bio | CAAGGGTAGCAACGGCTACAGAG TTT/3Bio/ | IDT |
| Handle site helix7, #7 -Bio | TTTAGGAACCCATGTACCGTAAC TTT/3Bio/ | IDT |
| Handle site helix7, #8 -Bio | ACGAGGTTGAGGCAGGTCAGACG TTT/3Bio/ | IDT |
| Handle site helix7, #9 -Bio | AAAAAATTCATATGGTTTACCAG TTT/3Bio/ | IDT |
| Handle site helix7, #10 | ACTATTATTTATCCCAATCCAAA | IDT |
| Handle site helix7, #11 | TAGTAAAGTAATTCTGTCCAGAC | IDT |
| Handle site helix7, #12 | TTAATCCTTGAAAACATAGCGAT | IDT |
| Handle site helix7, #13 -Bio | AGACCAGAAGGAGCGGAATTATC TTT/3Bio/ | IDT |
| Handle site helix7, #14 | GAATACGTGGCACAGACAATATTT | IDT |
| Handle site helix8, #0 | GCTTGCGTTGCGCTCACTGCCCG | IDT |
| Handle site helix8, #1 | CGGGCGCGGTTGCGGTATGAGCC | IDT |
| Handle site helix8, #2 | ACTGTTTACCAGTCCCGGAATTT | IDT |
| Handle site helix8, #3 | TTGTAGCCAGCTTTCATCAACAT | IDT |
| Handle site helix8, #4 | CATTTGCGGGAGAAGCCTTTATT | IDT |
| Handle site helix8, #5 | CGCGCGTTTTAATTCGAGCTTCA | IDT |
| Handle site helix8, #6 | GAGGCTCATTATACCAGTCAGGA | IDT |
| Handle site helix8, #7 | TTATACGTAATGCCACTACGAAG | IDT |
| Handle site helix8, #8 | ATGCCCTCATAGTTAGCGTAACG | IDT |
| Handle site helix8, #9 | CCAGCGCAGTCTCTGAATTTACC | IDT |
| Handle site helix8, #10 | AATTGACGGAAATTATTCATTAA | IDT |
| Handle site helix8, #11 | AAAGAGAATAACATAAAAACAGG | IDT |
| Handle site helix8, #12 | GTAATAGATAAGTCCTGAACAAG | IDT |
| Handle site helix8, #13 | CCGGTCTGAGAGACTACCTTTTT | IDT |
| Handle site helix8, #14 | CATTTGGATTATACTTCTGAATA | IDT |
| Handle site helix11, #0 | CTCTCACCAGTGAGACGGG | IDT |
| Handle site helix11, #1 | AAAGGCCCTGAGAGAGTTG | IDT |
| Handle site helix11, #2 | ACGCCAGCAGGCGAAAATC | IDT |
| Handle site helix11, #3 | GCACAAAATCCCTTATAAA | IDT |
| Handle site helix11, #4 | AGTGAGTGTTGTTCCAGTT | IDT |
| Handle site helix11, #5 | ACACGTGGACTCCAACGTC | IDT |
| Handle site helix11, #6 | CATCGATGGCCCACTACGT | IDT |
| Handle site helix11, #7 | TGTGGGGTCGAGGTGCCGT | IDT |

| Name | Sequence | Vendor |
|---|---|---|
| Handle site helix11, #8 | CTGGAGCCCCCGATTTAGA | IDT |
| Handle site helix11, #9 | GCCGGCGAGAAAGGAAGGG | IDT |
| Handle site helix11, #10 | CAGGGCGCTGGCAAGTGTA | IDT |
| Handle site helix11, #11 | AATACCCGCCGCGCTTAAT | IDT |
| Handle site helix11, #12 | TGCTTGCTTTGACGAGCAC | IDT |
| Handle site helix11, #13 | ACACAGAGCGGGAGCTAAA | IDT |
| Handle site helix11, #14 | CCGACAGGAACGGTACGCC | IDT |

**Supplementary Table 2 | Sequences for the three-color DNA-origami nanoruler.**

| | |
|---|---|
| **Imaging Parameters** | |
| Photon conversion factor | 1.84 |
| Linear (EM) gain | 1.0 |
| Pixel size [nm] | 159.0 |
| Time interval [ms] | 110 |
| Z-step [nm] | 50.0 |
| Camera offset [electron counts] | 91.0 |
| Read noise [electron counts] | 9.84 |
| **Find Maxima** | |
| Pre-Filter | None |
| Noise tolerance | 100 (TetraSpeck™ beads)<br>50 (Three-color dynein)<br>100 (Three-color nanoruler) |
| **Fit Parameters** | |
| Dimensions | 1 |
| Filter | Simplex-MLE |
| Max Iterations | 500 |
| Box size [pixel] | 6.0 |
| Fix width | Not selected |
| **Filter Data** | Nothing selected |
| **Positions** | Only imaged at one position except for TetraSpeck beads™ which were imaged as previously described[1] |
| **Skip Channels** | Not selected |

**Supplementary Table 3 | Fitting parameters used in µManager's 'Localization Microscopy' plug-in[1].**
