## Supplementary Movies for "Three-color single-molecule imaging reveals conformational dynamics of dynein undergoing motility": Supplementary-Movie_Captions.pdf

**Supplementary Movie 1**

The movie shows two examples of the movement of a three-color dynein along a microtubule. Note that the microtubule is not visible. The microtubule-binding domain (MTBD) of each motor domain and one of the two AAA rings are labeled with FluoroCubes. For one motor domain the AAA ring is labeled with a six dye ATTO 488 FluoroCube (green) and the MTBD is labeled with a six dye ATTO 647N FluoroCube (red). For the other motor domain only the MTBD is labeled with a six dye Cy3N FluoroCube (blue). The time is given in min:sec. Scale bar is 1 um.

**Supplementary Movie 2**

The movie shows an example of the step tracking of a three-color dynein while moving along a microtubule. Dynein was labeled as indicated in the cartoon on the top left. On the top right, we show a total internal reflection fluorescence (TIRF) microscopy movie of a three-color dynein moving along a microtubule. Note that the microtubule is not visible. On the bottom are the tracked dwells of the three labeled domains from the TIRF microscopy movie in x-and y-space. The time is given in min:sec. Scale bar is 500 nm.

**Supplementary Movie 3**

Example movie of a Monte Carlo simulation of dynein stepping along a microtubule in which all requirements / biases (**Fig. 6, Supplementary Fig. 6-8**) were applied (wild-type). One motor domain is represented by a cyan AAA ring and a blue MTBD while the other motor domain is represented by an orange AAA ring and a red MTBD. Note that there is no tail linking both motor domains together and therewith no physical constraint holding both motor domains together. For the simulation we used a 13 protofilament microtubule. Here, each parallelogram represents a tubulin dimer consisting of one copy of $\alpha$ and $\beta$ tubulin. Since dynein is able to rotate around microtubules, we added four additional lattices (opaque) to allow for right-handed and left-handed rotations.

**Supplementary Movie 4**

Example movie of a Monte Carlo simulation of dynein stepping along a microtubule in which the bias to step forward was ignored (**Supplementary Fig. 8**). One motor domain is represented by a cyan AAA ring and a blue MTBD while the other motor domain is represented by an orange AAA ring and a red MTBD. Note that there is no tail linking both motor domains together and therewith no physical constraint holding both motor domains together. For the simulation we used a 13 protofilament microtubule. Here, each parallelogram represents a tubulin dimer consisting of one copy of $\alpha$ and $\beta$ tubulin. Since dynein is able to rotate around microtubules, we added four additional lattices (opaque) to allow for right-handed and left-handed rotations.

**Supplementary Movie 5**

Example movie of a Monte Carlo simulation of dynein stepping along a microtubule in which the bias of the stepping direction along the off-axis was ignored (**Supplementary Fig. 8**). One motor domain is represented by a cyan AAA ring and a blue MTBD while the other motor domain is represented by an orange AAA ring and a red MTBD. Note that there is no tail linking both motor domains together and therewith no physical constraint holding both motor domains together. For the simulation we used a 13 protofilament microtubule. Here, each parallelogram represents a tubulin dimer consisting of one copy of $\alpha$ and $\beta$ tubulin. Since dynein is able to rotate around microtubules, we added four additional lattices (opaque) to allow for right-handed and left-handed rotations.

**Supplementary Movie 6**

Example movie of a Monte Carlo simulation of dynein stepping along a microtubule in which the bias of the stepping direction along the on-axis was ignored (**Supplementary Fig. 8**). One motor domain is represented by a cyan AAA ring and a blue MTBD while the other motor domain is represented by an orange AAA ring and a red MTBD. Note that there is no tail linking both motor domains together and therewith no physical constraint holding both motor domains together. For the simulation we used a 13 protofilament microtubule. Here, each parallelogram represents a tubulin dimer consisting of one copy of $\alpha$ and $\beta$ tubulin. Since dynein is able to rotate around microtubules, we added four additional lattices (opaque) to allow for right-handed and left-handed rotations.

**Supplementary Movie 7**

Example movie of a Monte Carlo simulation of dynein stepping along a microtubule in which the bias in stepping frequency of leading and trailing motor domain was ignored (**Supplementary Fig. 8**). One motor domain is represented by a cyan AAA ring and a blue MTBD while the other motor domain is represented by an orange AAA ring and a red MTBD. Note that there is no tail linking both motor domains together and therewith no physical constraint holding both motor domains together. For the simulation we used a 13 protofilament microtubule. Here, each parallelogram represents a tubulin dimer consisting of one copy of $\alpha$ and $\beta$ tubulin. Since dynein is able to rotate around microtubules, we added four additional lattices (opaque) to allow for right-handed and left-handed rotations.

**Supplementary Movie 8**

Example movie of a Monte Carlo simulation of dynein stepping along a microtubule for which no bias towards alternating stepping is enforced (**Supplementary Fig. 8**). One motor domain is represented by a cyan AAA ring and a blue MTBD while the other motor domain is represented by an orange AAA ring and a red MTBD. Note that there is no tail linking both motor domains together and therewith no physical constraint holding both motor domains together. For the simulation we used a 13 protofilament microtubule. Here, each parallelogram represents a tubulin dimer consisting of one copy of $\alpha$ and $\beta$ tubulin. Since dynein is able to rotate around microtubules, we added four additional lattices (opaque) to allow for right-handed and left-handed rotations.

**Supplementary Movie 9**

Example movie of a Monte Carlo simulation of dynein stepping along a microtubule for which no relative movement between AAA ring and MTBD is allowed (**Supplementary Fig. 6, 8**). One motor domain is represented by a cyan AAA ring and a blue MTBD while the other motor domain is represented by an orange AAA ring and a red MTBD. Note that there is no tail linking both motor domains together and therewith no physical constraint holding both motor domains together. For the simulation we used a 13 protofilament microtubule. Here, each parallelogram represents a tubulin dimer consisting of one copy of $\alpha$ and $\beta$ tubulin. Since dynein is able to rotate around microtubules, we added four additional lattices (opaque) to allow for right-handed and left-handed rotations.

**Supplementary Movie 10**

Example movie of a Monte Carlo simulation of dynein stepping along a microtubule in which only inter-MTBD distance independent movement between AAA ring and MTBD is allowed (**Supplementary Fig. 6, 8**). One motor domain is represented by a cyan AAA ring and a blue MTBD while the other motor domain is represented by an orange AAA ring and a red MTBD. Note that there is no tail linking both motor domains together and therewith no physical constraint holding both motor domains together. For the simulation we used a 13 protofilament microtubule. Here, each parallelogram represents a tubulin dimer consisting of one copy of $\alpha$ and $\beta$ tubulin. Since dynein is able to rotate around microtubules, we added four additional lattices (opaque) to allow for right-handed and left-handed rotations.
